# Generative continuous time model reveals epistatic signatures in protein evolution

**DOI:** 10.1101/2025.09.17.676821

**Authors:** Andrea Pagnani, Pierre Barrat-Charlaix

**Affiliations:** DISAT, Politecnico di Torino, Corso Duca degli Abruzzi 24, 10129, Torino, Italy; Italian Institute for Genomic Medicine, IRCCS Candiolo, SP-142, 10060, Candiolo, Italy; INFN, Sezione di Torino, Via Pietro Giuria 1, 10125 Torino, Italy; DISAT, Politecnico di Torino, Corso Duca degli Abruzzi, 10129, Torino, Italy; Sorbonne Université, CNRS, Institut de Biologie Paris-Seine, Laboratoire de Biologie Computationnelle, Quantitative et Synthétique, Paris F-75005, France

**Author notes:** Correspondance to: PBC.

**Keywords:** sequence evolution model, generative model, epistasis, protein evolution

## Abstract

Protein evolution is fundamentally shaped by epistasis, where the effect of a mutation depends on the sequence context. As standard phylogenetic methods assume independently evolving sites, there is a need for more complex models based on accurate estimations of the fitness landscape. Good candidates are modern generative models – such as the Potts model – which successfully capture epistatic effects. However, recent works on generative evolutionary models usually use discrete time, making them difficult to integrate with the standard frameworks in evolutionary biology. We introduce a continuous-time sequence evolution model using the Gillespie algorithm and parameterized by a generative Potts model. This approach enables us to simulate realistic, family-specific evolutionary trajectories and allows for direct comparison with independent-site models. Surprisingly, we find that while epistasis significantly slows down evolution, it does not change the average evolutionary rates at individual sites. This is explained by the rate heterogeneity caused by context-dependence: we show that the rate at some positions varies between null to high values depending on the context, while other positions are essentially independent from the context. Finally, we show that epistasis leads to a systematic underestimation bias in the inference of evolutionary distance between sequences. Overall, our work provides a new tool for simulating realistic protein evolution and offers novel insights into the complex interplay between epistasis and evolutionary dynamics.

**Significance statement:** Understanding how proteins evolve is central to molecular biology and phylogenetics. Traditional evolutionary models assume that mutations act independently at each position in a sequence. This neglects epistasis — the fact that the effect of a mutation depends on the rest of the sequence — which is known to be ubiquitous in proteins. By simulating protein evolution in continuous time using a generative model, our approach produces realistic sequences and reveals how epistasis shapes evolutionary dynamics. We find that epistasis slows down evolution and can mislead common methods for estimating evolutionary timescales. This work bridges modern generative models of proteins and phylogenetics, providing new tools to better understand molecular evolution.

## I. INTRODUCTION

Proteins are essential molecules for all forms of life. They fill very specific functions that evolution has optimized during billions of years. For this reason, members of a protein family that share a common biological function can have sequences differing in up to 80% of their amino acids. On the other hand, a handful of ill-chosen mutations can render a protein completely non-functional. Sequence evolution models are theoretical tools that describe the evolutionary process by quantifying how protein sequences change with time. They are the central element of phylogenetic inference, which aims at reconstructing the evolutionary relations between genes and organisms.

A large amount of work has been dedicated to making sequence evolution models more realistic. This includes modeling the heterogeneity of evolutionary rates at different sequence positions [1, 2], the construction of matrices quantifying the transition rates between amino acids [3, 4], or the development of more complex methods such as site-specific amino acid frequencies [5–7] or heterotachous rates [8]. Virtually all evolutionary models make the simplifying assumption that different positions in the protein sequence evolve independently. However, there is little doubt that sites in a protein sequence are not independent and that this influences evolution [9, 10]. This interdependence is called epistasis, and neglecting it is in fact a known weakness of evolutionary models [11].

The last decade has seen important progress in the development of generative protein sequence models. These models assign a probability to protein sequences that can be sampled from, with recent implementations being capable of generating sequences that are functional or at least statistically indistinguishable from natural ones [12, 13]. This probability distribution is learnt by taking advantage of the fact that many homologous proteins share similar biological functions but can have very different sequences. Generative sequence models can take the form of restricted Boltzmann machines [14], variational auto-encoders [15], or deep neural networks [16, 17]. A particularly interesting class of models is the Potts model, because of both its mathematical simplicity and the possibility to interpret its parameters biologically [18]. Potts models were first developed to predict structural contacts in the protein structure [19, 20] and were later found to generate sequence that closely match the natural ones if inferred with the proper algorithm [21]. It must be stressed that the key ingredient in all these techniques is the accurate modeling of epistasis: the fact that the effect of a mutation in a protein sequence depends on the context, that is on the rest of the sequence.

Integrating epistasis into sequence evolution models with the perspective of phylogenetic inference is a challenging problem and has been addressed in the past. In [22, 23], authors design a method to identify pairs of epistatic positions and to reconstruct the phylogenetic tree, while [24] shows that the inclusion of epistasis typically degrades phylogenetic reconstruction done by a traditional site-independent evolutionary model. However, these works are based on the assumption of non-overlapping epistatic pairs, which can be considered a strong oversimplification given the state of the art on generative models. More general models of epistasis have also been used: in [25, 26], protein fitness is assumed to be a sum of pairwise interactions corresponding to contacts in the tertiary structure, with values given by a biochemistry-based statistical potential [27]. Authors then showed that it is possible to infer certain parameters of the evolutionary model given sequence data. Using the same idea, [28] shows that the inclusion of epistasis typically degrades phylogenetic reconstruction. While this approach is closer in spirit to modern sequence models, using a biochemistry based potential and restricting interactions to contacts makes it very unlikely that the resulting model generates functional sequences.

Recently another line of research has built on modern generative models to understand the consequences of epistasis on evolutionary dynamics. In [29], authors show that many effects that need to be explicitly modeled by independent-sites evolutionary models, such as Gamma distributed rates, heterotachy or “Stokes shift” [30] emerge naturally if an epistatic model is used. Epistasis has also been shown to introduce different timescales in evolution: fast mutational processes within a sequence context, and slow change of the context itself [31]. Generative evolutionary models can also be used to model the results of directed evolution experiments, or to evolve new functional sequences starting from a natural one [32, 33]. Potts-like kinetic simulations have also been applied to study the ordering and timing of mutations under directional selection pressure, for instance to model the emergence of drug resistance [34, 35]. These works are based on discrete Monte-Carlo sampling of a Potts model. The qualitative picture of epistatic effects on evolution is expected to be consistent between discrete and continuous approaches, as indeed found when comparing our results to prior work [29, 31]. Discrete Monte Carlo is very natural, but is at odds with traditional evolutionary models used in phylogenetics which are based on a continuous time approach; the use of continuous time enables quantitative connections that are not accessible with discrete-time models. First, the parameter *t* in a continuous model corresponds to on average one substitution per sequence position — the same unit as phylogenetic branch lengths — allowing direct comparison with evolutionary distances inferred by phylogenetic reconstruction. Secondly, as we will show in the results, using continuous time makes it possible to discuss substitution rates at the sequence and site levels and to compare them directly to those of site-independent models.

In this article, we propose a sequence evolution model that is generative and also compares more naturally to the existing literature. We base it on a Potts model with a continuous time evolution procedure based on the Gillespie algorithm [36]. Potts models inferred using the bmDCA algorithm are accurate generative models [12, 13], and sequences obtained during an evolutionary trajectory will therefore be statistically very similar to natural ones and likely functional. Potts models are learned using single families of homologous proteins and are thus family specific, but our simulation procedure is general: our model can simulate evolution for any protein family at the cost of inferring a Potts model. We propose this model as an open source package that can be used by the community (Code Availability section).

We then set on using our model to study the consequences of epistasis on protein evolution. In particular we observe that epistasis slows down evolution, and yet the average substitution rates at both the sequence and position level are very close to the ones of a site-independent model. This is explained by the fact that the epistatic model evolves at different rates in different contexts. We then derive a method to score sequence positions by how context-dependent their evolution is. Finally, we show that the inference of the evolutionary time between two sequences is strongly biased when epistasis is present, and that this is caused by the existence of positions whose evolution is strongly context-dependent.

## II. RESULTS

### A. Continuous time generative evolution model

A Potts model is defined at the level of a protein family and associates a probability to each aligned amino acid sequence **a** = {*a*_1_ … *a*_*L*_} of fixed length *L*:

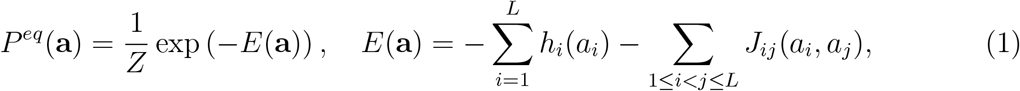

where *Z* is a normalization constant, parameters *h* and *J* are called fields and couplings and the label “eq” stands for “equilibrium”, as explained below. The state *a*_*i*_ at position *i* of **a** can take *q* = 21 values: the 20 natural amino acids and the gap symbol. Potts models have been shown to be accurate generative models in the sense that the distribution *P*^*eq*^ closely matches that of natural sequences, and that sequences sampled from *P*^*eq*^ are generally found to be functional [12, 33]. Parameters *h* and *J* are calculated to fit the first and second order marginals *π*_*i*_(*a*) and *π*_*ij*_(*a, b*) of *P*^*eq*^ to the single-site and pairwise distribution of amino acids found in the alignment of natural sequences.

We want to construct a sequence evolution model with two properties. First, it should be parametrized using continuous time, similarly to most models used in phylogenetics. In practice, this means that the model is a continuous time Markov chain (CTMC) characterized by a sequence-to-sequence transition rate matrix *Q* of size *q*^*L*^ × *q*^*L*^. While this matrix is of gigantic dimensions, we show below that it never needs to be fully computed. Secondly, we want *P*^*eq*^ to be a fixed point of the dynamics: whatever the starting sequence, sampling the model after a long time *t* should be equivalent to sampling from the generative distribution. These two properties can be expressed mathematically as

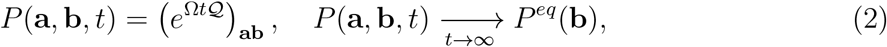

where *P* (**a, b**, *t*) is the transition probability from sequence **a** to **b** in evolutionary time *t*. Following a common practice in phylogenetics, we scale the time parameter *t* by a model-dependent factor Ω so that the average rate of substitutions per position is one. Ω can be computed using Monte-Carlo sampling of *P*^*eq*^ (Material and Methods).

To ensure that *P*^*eq*^ is a fixed point of the dynamic, we require that the transition rate matrix verifies the detailed balance equations: for any pairs of sequences **a, b** we want *P*^*eq*^(**a**)*Q*_**ab**_ = *P*^*eq*^(**b**)*Q*_**ba**_. In this work, we enforce this by using the Glauber dynamics defined by

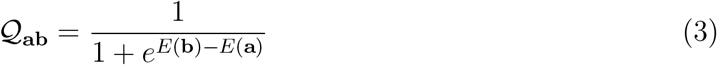

with *E* the energy function of the Potts model. Other dynamics are possible, such as Metropolis, Gibbs, square-root or mutation-selection (see Supplementary Section A 1); the latter is of particular interest as it has a direct interpretation in terms of population genetics and has been used previously in the context of phylogenetics [5, 26]. However, our main results do not qualitatively depend on this choice and find dynamics other than Gibbs to be quantitatively very similar to Glauber. Note that unlike Metropolis-style kinetic Monte Carlo, where simulation steps correspond to proposals that can be accepted or rejected, the Gillespie algorithm works using the average number of substitutions per site *t*.

To further simplify the model, we only consider the case where *Q*_**ab**_ is non-zero only if **a** and **b** differ by only one mutation. This amounts to saying that only single amino acid mutations have a non vanishing probability for very short evolutionary times. This simplification is essential to simulating dynamics using Gillespie’s algorithm [36], as in this case the rows of the Q matrix only have *L*(*q* − 1) non-zero terms. If we note N(**a**) as the set of sequences that are one mutation away from **a**, we can easily compute the rate of substitution starting from sequence **a** as

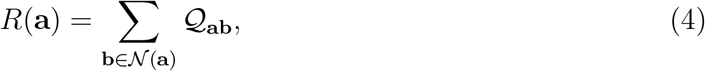

Computing this sum is numerically tractable as it only involves *L*(*q* − 1) terms.

This modeling strategy is well-suited for a Potts model but can in principle be applied to any distribution *P*^*eq*^. We can therefore use it to simulate evolution under a profile model, which considers sequence positions as independent but reproduces the frequencies of amino acids in each column of the alignment. Formally, the equilibrium distribution for the profile model is the same as the one of Eq. 1 but with coupling parameters set to zero. The fields **h** are adjusted so that the frequency of amino acids at each position remains the same, see Eq. 10 in Material and Methods. We show in Supplementary Section A 2 that in this case the transition probabilities of Eq. 2 take a factorized form with a position-specific average substitution rate. The essential difference between a Potts and a profile model is therefore the absence of epistasis in the latter.

In the following, we explore properties of our evolutionary model using a specific protein family (PF00072, response regulator). We use its Pfam alignment [37] to learn a Potts model and a profile model, allowing us to explore the effect of epistasis on evolution. Additional results for two other families are shown in the Supplementary Material (PF00595, PDZ domain; PF13354, beta-lactamase). We learn the Potts models using the bmDCA algorithm [38] (Material and Methods).

### B. Generative properties

In Supplementary Figure 1, we show that our generative model is accurate in a statistical sense. To do so, we start evolutionary trajectories from *M* = 1000 different natural sequences of the PF00072 family. Each trajectory is sampled at times [0.1, 0.5, 1, 5, 20], so that we obtain an alignment of *M* artificially evolved sequences at each of these times. We then compare the statistical properties of the natural and evolved sequences. Specifically, we plot the single-site frequencies and pairwise correlation of amino acids in the alignments; the projection of generated sequences onto the principal components of the natural alignment; the distribution of the statistical energy score Eq. 1. The figure clearly shows that our artificially evolved sequences are statistically very close to the natural ones. This result is expected given Eq. 2 and the way parameters *J* and *h* are fitted.

**FIG. 1.**
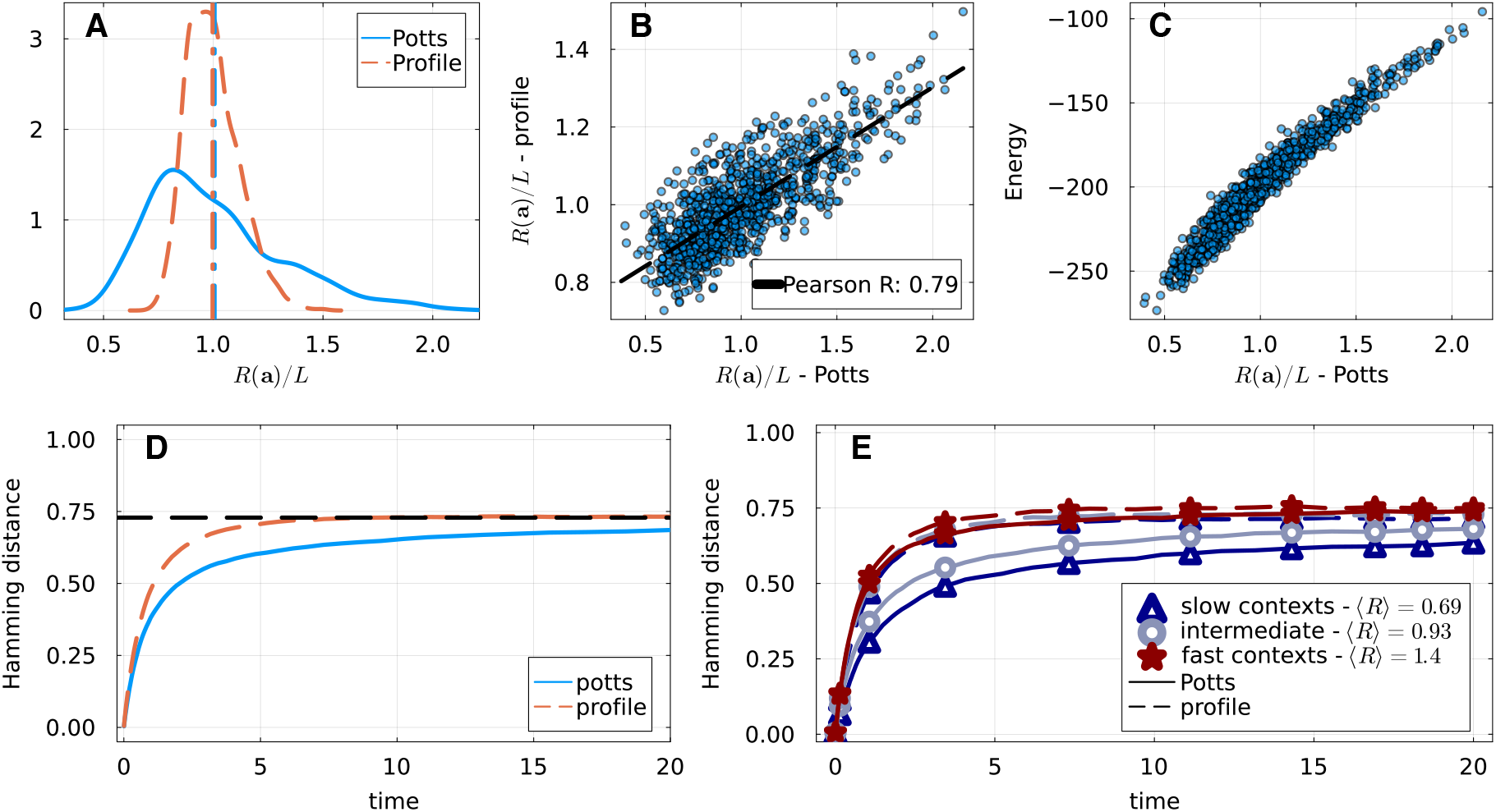
Rate of evolution, for models fitted to the PF00072 family. **A**: Distribution of the scaled sequence rates *R*(**a**) over natural sequences for the profile and Potts models. Rates are scaled by the length of the sequence *L*, so that they are one on average. **B**: Comparison of rates in the profile and Potts model over natural sequences, showing a high correlation. **C**: Comparison of sequence rate and energy in the Potts model over natural sequences. Sequences with low energy, that is more probable, accumulate fewer substitutions in a given amount of time. **D**: Average sequence divergence versus time for evolution simulated with a Potts and profile models. The average is over a set of 100 starting sequences and 5 realizations for each starting sequence. Sequence divergence is computed relative to the initial sequence. The dashed horizontal line shows the average distance between natural sequences and the initial sequences used here. **E**: Average sequence divergence versus time segregated by the category of the starting sequence: slow ([0, 0.33] quantile), intermediate ([0.34, 0.66] quantile) and fast ([0.67, 1.] quantile). The scaled average rate *R*(**a**) of initial sequences in each category is indicated in the legend.

The inclusion of the coupling terms modeling epistasis is a key ingredient in this matter. Supplementary Figure 2 shows the same experiment using a profile model. In this case, the single site distribution of amino acids is maintained at long evolutionary times, but the other metrics show a clear difference between natural and evolved sequences at intermediate and long times (*t >* 1). We also include results obtained for the LG+G4 model which we simulated using the alisim software [4, 39] in Supplementary Figure 3. As this type of model cannot capture the amino acid frequencies of the natural alignment, its generative properties are worse than those of the profile model. In both cases, the values of the statistical energy increase rapidly with time, leaving the range of energy of natural sequences. This is relevant as the values of the statistical energy were found in [12] to be somewhat predictive of *in vivo* functionality: sequences with an energy value higher that the typical energy of the natural sequences were most of the time not functional.

**FIG. 2.**
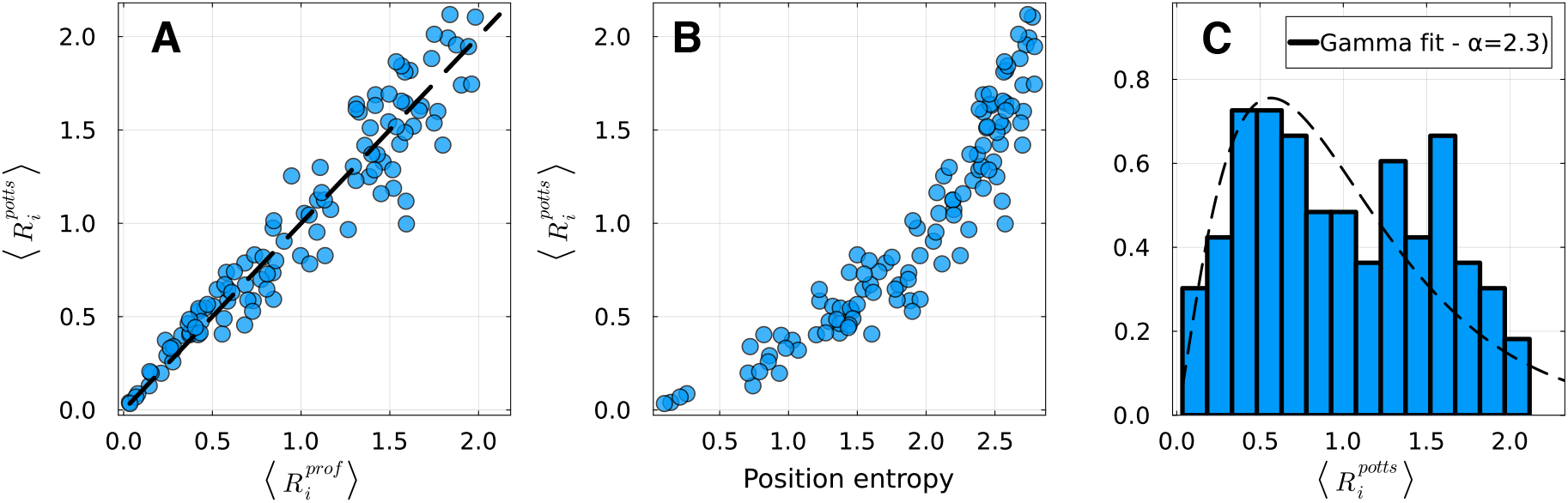
**A**: average evolutionary rate ⟨*R*_*i*_⟩ at site *i* for the Potts and the profile models. The average is performed over natural sequences. **B**: average rate ⟨*R*_*i*_⟩ against the entropy of position *i* in the alignment. More conserved positions accumulate fewer substitutions per unit of time. **C**: distribution of average site-specific rates over sites *i*. The dashed line shows a maximum-likelihood fit to a Gamma distribution (parameters *α* = *β* = 2.3).

**FIG. 3.**
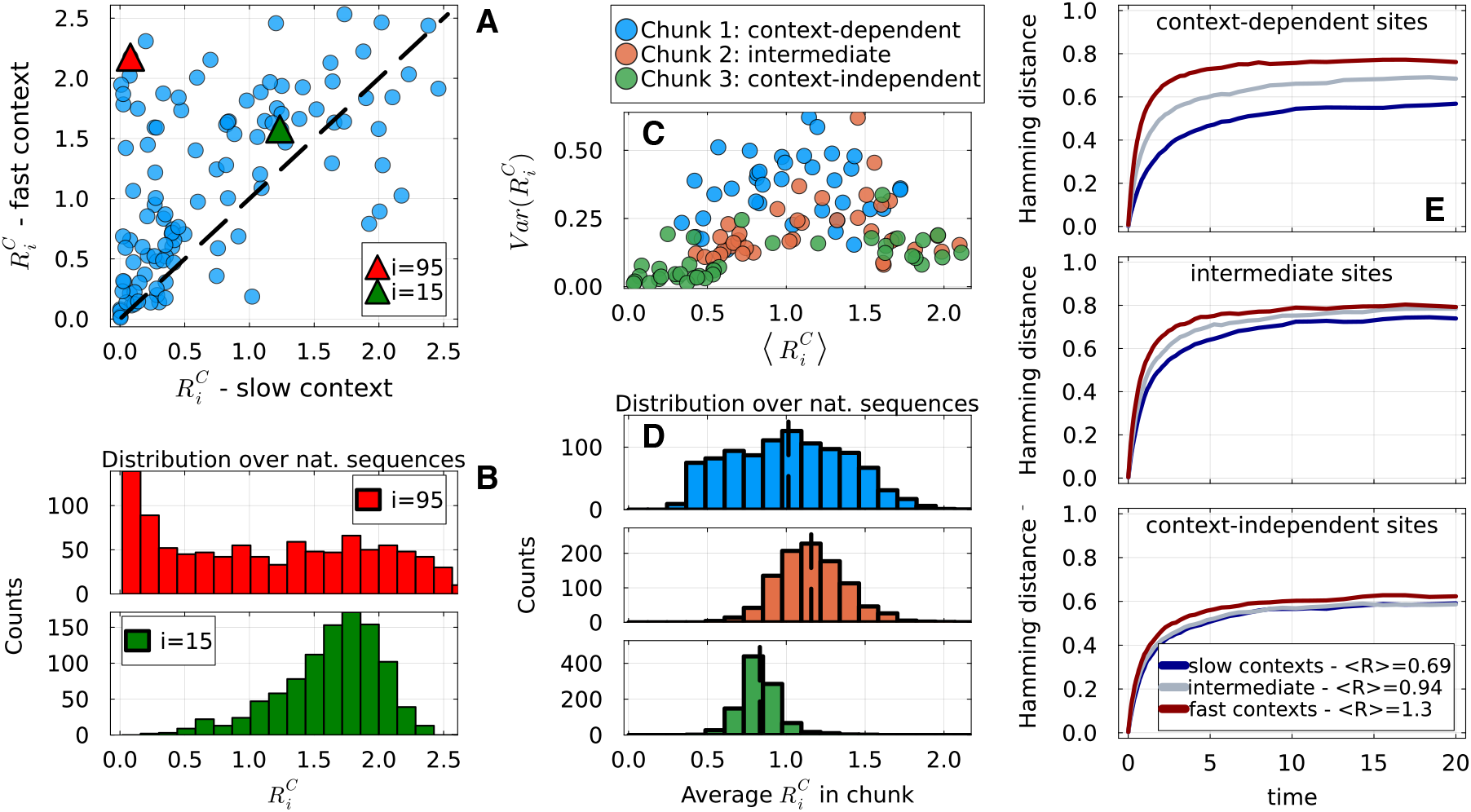
**A**: context-dependent rates at each site for a slow and a fast context (see panel **E** of Figure 1). **B**: Distribution of context-dependent rates across natural sequences at the two highlighted positions. **C**: Variance versus mean of 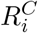 across natural sequences, for each position *i*. The color code corresponds to the division of sites in three categories with different importance of the context. **D**: Distribution of the average 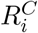 inside each category across natural sequences. Colors are the same as in panel **C. E**: Evolutionary trajectories showing the Hamming distance to the starting sequence as a function of time, for three starting sequences (slow, intermediate, fast contexts). Each plot focuses on one category, with Hamming distance being computed using only the corresponding positions.

### C. Sequence rates, epistasis and speed of evolution

Our model assigns a rate of evolution *R*(**a**) to each sequence **a**, as defined in Eq. 4. This sequence specific rate determines how fast evolution proceeds when starting from **a**: it is precisely equal to the inverse of the average waiting time to the next substitution (Material and Methods). In Figure 1**A&B** (Supplementary Figure 4 & Supplementary Figure 5 for extra families), we show the distribution of *R*(**a**)*/L* for sequences coming from the alignment of the PF00072 family and for both the profile and the Potts models. Scaling by the sequence length *L* allows us to have values that are on average one for both models. Figure 1**A** shows that the distributions of *R*(**a**) have sensibly different width for the two models, with the Potts giving rise to more varied rates (standard deviation 0.29) than the profile (standard deviation 0.12). Consequently, the number of substitutions as a function of time varies more strongly for the case of the Potts model Supplementary Figure 6. On the other hand, rates in the profile and Potts models are quite correlated as is shown on Figure 1**B**, with a linear correlation of about 0.8. Neglecting epistasis thus tends to uniformize the rates of evolution without affecting which sequences have high or low rates. It is interesting to note that since different sequences have different rates, our model will by design feature heterotachy [40], that is the change in evolutionary rates along lineages. Indeed, *R*(**a**) changes during the evolutionary trajectory as **a** itself changes. As the figure shows, this is not an intrinsic property of epistasis. However, epistasis makes the *R*(**a**) distribution wider, so we expect heterotachy to be more pronounced in this case, consistently with [29].

**FIG. 4.**
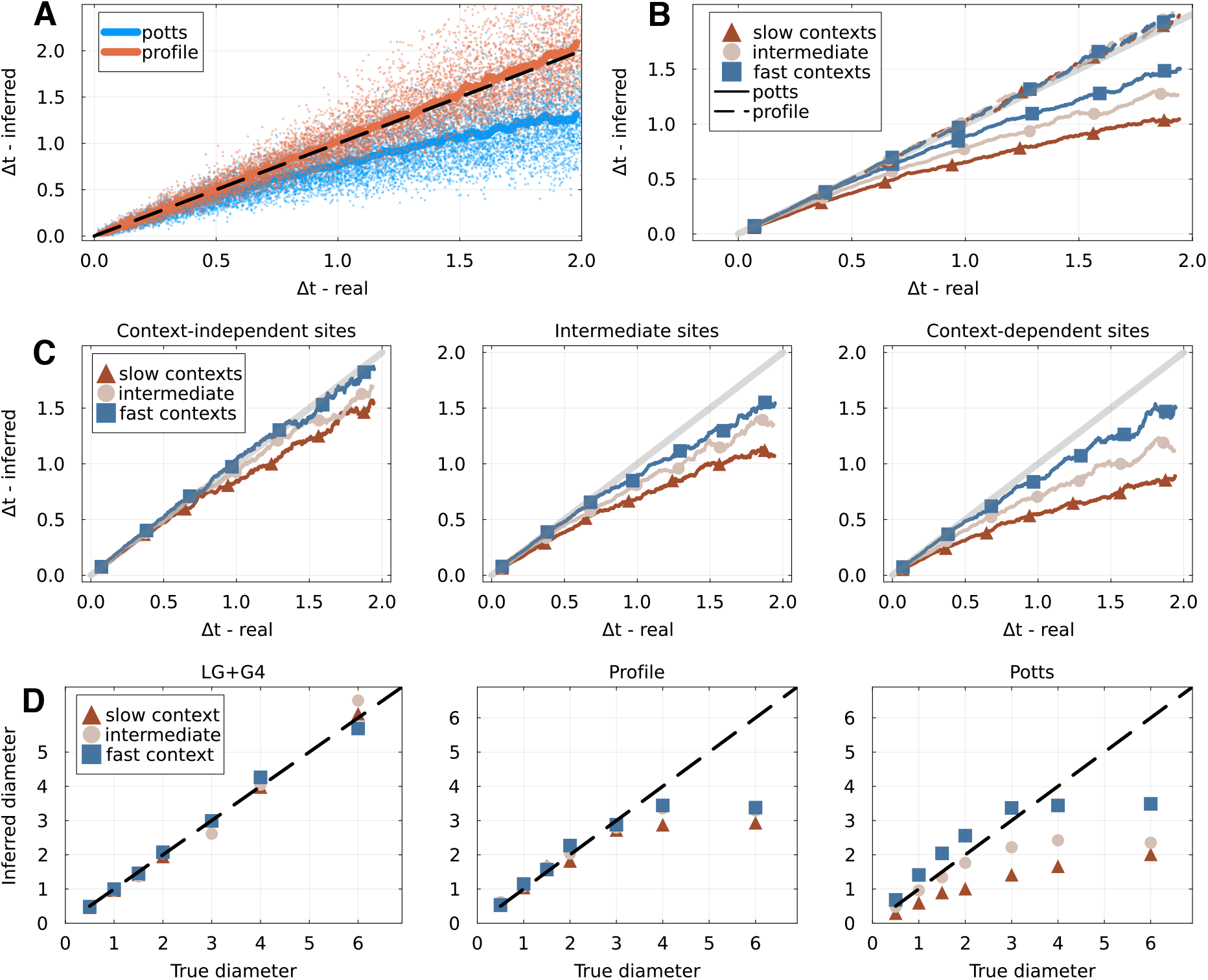
**A**: reconstructed versus real evolutionary distance for all simulated pairs of sequences, for the profile and Potts models. One point represents a pair of sequences, and the smooth lines are obtained using a running average. **B**: reconstruction of time *t* for different contexts: three groups of pairs of sequences are made, based on the evolutionary rate of the starting sequence. Lines show the average reconstructed versus real time for each group. The Potts model is shown as a solid line, the profile as a dashed line. **C**: reconstruction of time *t* when focusing on specific parts of the sequences, as defined in the previous section: the context-independent (left), intermediate (center) and context-dependent sites. **D**: inference of the tree diameter in the four-taxon experiments, with three different evolutionary models (Alisim, profile and Potts). Different initial sequences (slow, intermediate and fast contexts) are indicated by different colors.

Another effect of epistasis that has already been observed is to slow down evolution [29, 31]. We confirm this by running evolutionary trajectories starting from a set of 100 natural sequences of the PF00072 family for a total time *t* = 20. For each starting sequence, we run 5 independent trajectories, and monitor the Hamming distance between the sequence at time *t* and the initial one. Figure 1**D** shows the average divergence as a function of time obtained in this process. We observe that evolution under the Potts model is on average slower than under the profile model in the sense that it takes a longer time, or equivalently a higher number of substitutions, to reach a given Hamming distance. Since both models reproduce the distribution of amino acids in each column of the alignment, they have the same average sequence divergence on the long-term. It is important to note that because of our scaling, the meaning of time for the two models is the same: in time interval one, on average *L* amino acid substitutions occur. This can be verified in Supplementary Figure 1**C**, where sequence divergence is plotted against the number of substitutions. The fact that it takes a longer time to reach a given Hamming distance from the initial sequence under the epistatic Potts model indicates that substitutions occur at a more restricted set of sequence positions or involve more reverse mutations.

A related question is what determines the rate of a particular sequence. In Figure 1**C**, we show that the sequence-specific rate is highly correlated with the energy *E*(**a**) of the sequence in the Potts model. This is in fact expected, as for all dynamics but Gibbs, the substitution rate between two sequences depends monotonously on their energy difference Δ*E*. It is likely that sequences with low energy are closer to a minimum, so that single site mutations have on average a higher Δ*E*, and hence correspond to a low substitution rate. This can also be interpreted biologically if one assumes that the energy of the Potts model is a good predictor of protein functionality [12]. Indeed, we expect highly optimized and functional proteins to be close to a fitness peak and therefore have few accessible non-deleterious substitutions, while high energy sequences correspond to a fitness valley and can more easily mutate. In fact, our sequence rate also correlates strongly with other measures of how fast a sequence evolves: we show for example in Supplementary Figure 7 that it is related to the context-dependent entropy as defined in [31]. Rates for alternative dynamics are also very correlated with the ones for Glauber, as shown in Supplementary Figure 8.

An advantage of *R*(**a**) as we define it is that it is directly interpretable in terms of the evolutionary process, as it is the inverse of the average waiting time to the next substitution. However, it is also a local quantity, in the sense that the rate of the sequence coming after **a** in the evolutionary trajectory could in principle be completely different. Therefore, an important question is whether this rate matters for long-term evolution. To evaluate this, we start evolutionary trajectories from initial sequences chosen from the quantiles ([0, 1*/*3], [1*/*3, 2*/*3], [2*/*3, 1]) of the distribution of Figure 1 **A**, which we call “slow”, “intermediate” and “fast” contexts in the following. As can be seen in Figure 1**E**, distance from the starting sequence increases more slowly for starting sequences with a low rate. This not only concerns the initial slope of the distance versus time curve but impacts the trajectory on the long-term: at *t* = 20, the trajectory starting from fast initial sequences is fully equilibrated while the one starting from the intermediate sequences has barely reached saturation and the one starting from the slow sequences is still far from saturation. The saturation values reached at infinite time can be derived analytically from single-site conservation and are different for these three trajectories, as the average distance of natural sequences to each particular starting sequence may differ. The profile trajectories (dashed lines) converge faster in all cases and indicate the value of saturation. Interestingly, the trajectory starting from the “fast” sequence evolves at the same speed in the profile and Potts model, indicating that epistasis only weakly influences evolution in this case.

Note that these results illustrate two effects of epistasis on evolutionary dynamics that are important to distinguish. First, interactions tend to increase the variance in sequence substitution rates, creating what we call slow and fast contexts. The number of substitutions per unit time in a given low-rate sequence is lower in the case of the Potts model than in the profile model (Figure 1**A**, Supplementary Figure 6**A&B**) It is this effect that we refer to when qualifying initial sequences using the adjectives “slow” and “fast”. The second effect is that for a given number of substitutions, epistasis slows down evolution, as illustrated by Figure 1**A&B** and Supplementary Figure 6**C&D**. This effect is shown to be much stronger for sequences with a low rate, as is visible in Supplementary Figure 6**D**: it takes about two substitutions per site for trajectories starting from fast contexts to reach Hamming distance ~ 0.6, but trajectories starting from slow contexts barely reach this value after ten substitutions per site. The two effects combine when measuring quantities as a function of time as in Figure 1.

### D. Average site-specific evolutionary rate

For a given sequence **a** and a position *i*, the site-specific rate *R*_*i*_(**a**) is defined as the sum of the rates from **a** to any mutant at position *i*. Defining *N*_*i*_(**a**) as the set of sequences that are mutants of **a** at position *i*, we have

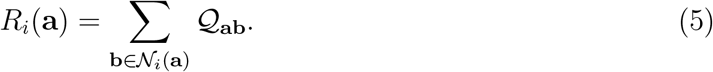

In order to study a quantity that does not depend on a particular sequence, we define the *average* site-specific rate ⟨*R*_*i*_⟩ as the average of *R*_*i*_ over natural sequences. ⟨*R*_*i*_⟩ represents the average rate at which substitutions occur at site *i* during simulated evolution.

A first question is whether epistasis influences the rates at individual sites *i*. Figure 2**A** (Supplementary Figure 9 & Supplementary Figure 10 for extra families) shows that this is not the case, as the values of ⟨*R*_*i*_⟩ between a Potts and a profile model are highly similar. Therefore, on average, epistasis does not slow down the substitution process at any site in particular. Figure 2**B** reveals that average substitution rate at a site is heavily correlated with conservation: positions that are variable in the alignment tend to evolve fast in the model, while conserved positions have a low rate. This is not surprising, and analytical calculations in the case of a profile model confirm that the average rate can always be linked to conservation (Supplementary Section A 3). The Potts and profile models both capture the conservation of columns in the alignment, explaining that sites evolve on average at the same rate in both cases. On the other hand, this does not explain what makes the epistatic dynamics different: we will show that it is necessary to consider rates in particular contexts for this purpose.

Another relevant question is the distribution of site-specific substitution rates. The importance of taking into account and correctly modeling the rate heterogeneity among sites is a well established principle in phylogenetics. Rates are usually modeled using categories that are based on a Gamma [1, 2] or parameter-free distribution [41]. In our case, the rate heterogeneity comes from the characteristics of the equilibrium distribution *P*^*eq*^: it can be computed exactly for the profile model (Eq. A7 of the Supplementary Material) and estimated numerically for the Potts. Figure 2**C** shows the distribution of ⟨*R*_*i*_⟩ for the Potts model. We observe that this distribution is compatible with a Gamma distribution of parameter *α* = 2.3 and mean one – by design, the mean of ⟨*R*_*i*_⟩ over *i* is one. However, this result should be taken with care as the quality of the fit appears lower for other dynamics, especially Gibbs dynamics, as can be seen in Supplementary Figure 11. Note that given the results of Figure 2**A**, this distribution would not change if we were to use the profile model. The differences in evolutionary dynamics caused by epistasis can therefore not be accounted for by simply modeling the average site rates.

The value of the shape parameter *α* = 2.3 reflects the variance of context-averaged rates ⟨*R*_*i*_⟩, which corresponds to the deep-phylogeny limit where sequences have explored many contexts along their branches; it is on the high-end of values reported for empirical protein alignments [4]. In a shallow phylogeny, the relevant quantity is instead the context-dependent rate 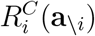 defined below in Eq. 6 – in a fixed ancestral context. Because epistasis increases the rate of some sites faster and decreases that of others depending on the background, the variance of 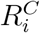 across sites in a given context exceeds the variance of ⟨*R*_*i*_⟩ and therefore corresponds to a lower *α*. This can be formalized using a variance decomposition, see Supplementary Section A 4. Supplementary Figure 12 shows that shape parameters inferred from rates in specific contexts are on the order of 1, closer to typical values. For the profile model, context-dependence vanishes and the two quantities coincide, predicting high value of *α* in any phylogeny.

### E. Context-dependent site evolutionary rate

In order to better understand the role of epistasis in the Potts dynamics, we want to quantify the dependence of the rate of evolution at each site on the sequence context. We define the *context-dependent rate* at a given position *i* as

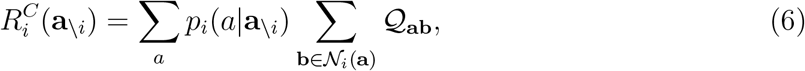

where **a**_*\i*_ designates a set of amino acids at positions other than *i, i*.*e*. a context, and *p*_*i*_(*a*|**a**_*\i*_) the conditional probability at position *i* given this context according to the generative model. Note that this conditional probability can be easily computed if a Potts model is used. The sequence **a** in the inner sum is defined as having context **a**_*\i*_ and state *a* at position *i*.

We expect 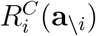 to be a marker of epistasis. Indeed, for a profile model with additive fitness, we always have 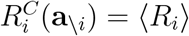 since the conditional probability *p*_*i*_ does not depend on the context. On the other hand, we can observe significant differences in rates for different contexts if we use the generative Potts model. Panel **A** of Figure 3 (Supplementary Figure 13 & Supplementary Figure 14 for extra families) shows the scatter plot of 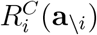 for the contexts of two particular sequences, at quantiles 10% (slow) and 90% (fast) of the distribution of *R*(**a**) (see Figure 1**A**). Many positions are found to have a very small rate in the slow context while having a significantly higher rate in the fast context. We choose two particular sites to highlight, with a procedure described further below: site 15 has roughly the same rate of about 1.5 in the two contexts, while site 95 evolves at a rate higher than 2 in the fast context and close to 0 in the slow context. In fact, Figure 3**B** shows the distribution of 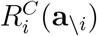 across all contexts found in the natural sequences and for the two highlighted positions *i* = 95 and *i* = 15. We find that the rates of site 15 are centered around an average value of 1.5 for all natural sequences, while the ones of site 95 are spread almost uniformly from 0 to 2.5. Similar analysis of other positions reveals that this is a common pattern: the evolutionary rate at some site depends weakly on the context, while others are very evolvable given certain contexts and “frozen” in others. The existence of such context-dependent sites is an effect of epistasis.

To explore this, we construct a ranking of sites based on how context-dependent their evolution is. We define 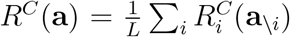 as the average rate of evolution of sites in the context defined by sequence **a**. We then rank sites by their contribution to the variance of *R*^*C*^(**a**) over sequences: the top *K* sites of the ranking are chosen so that the variance of *R*^*C*^(**a**) restricted to these *K* sites is maximal (see Material and Methods). This procedure makes the ranking depend both on the variance 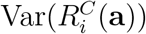 of the context-dependent rate over sequences and of the covariance 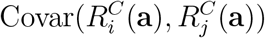 of context-dependent rate between positions. Our ranking therefore selects sets of positions that have a variable 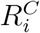 but also tend to be variable or frozen in similar contexts. Note that since our substitution rates are typically well correlated with other measures of mutability such as context-dependent entropy [32], a ranking based on the latter quantity would likely give comparable results. The effect of the ranking is visible on the heatmap of the covariance matrix 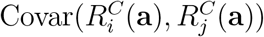 (Supplementary Figure 15). In Figure 3**A**, we highlighted the site ranked first and the one ranked at position 2*L/*3, with *L* the length of the sequences.

Interestingly, a visual inspection of structures suggests that the top-ranked sites have a structural interpretation in the PF00072 and PF00595 families. In both cases, the top *L/*5 sites – where *L* is the length of the sequence – are observed to be structurally close and located at a binding site or an interface, as is visible on Supplementary Figure 16. This interpretation is less evident in the case of the third family PF13354, where the top-ranked sites seem more spread-out over the structure. We note that this observation is qualitative and that a systematic structural analysis of how epistatic constraints emerge from structure is beyond the scope of this work.

Ideally, we would like to identify a set of positions whose evolution is context-dependent, and another that is context-independent. However, our ranking procedure does not provide a clear threshold to group sites. We therefore decide to divide positions into three categories of size *L/*3 based on the ranking; we call these categories context-dependent, intermediate and context-independent sites. Figure 3**C** shows for each *i* the variance against the average of 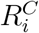, with a color corresponding to the category that the site belongs to. As expected, context-dependent sites tend to have a high variance and the context-independent sites a low variance. Interestingly, context-independent sites have either a high or a low average evolutionary rate, and not an intermediate one.

Figure 3**D** shows the distribution of the average 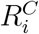 for *i* in a given category over contexts of natural sequences, that is the distribution of 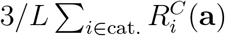. We observe a very wide distribution for context-dependent sites: this indicates the existence of contexts in which those sites jointly have a high or low substitution rate. On the other hand, the distribution for context-independent sites is very peaked, which is expected as they have a low individual variance. Note that for a profile model, there is no difference between the context-dependent rate and the average rate: we would then observe perfectly peaked distributions for all categories. Interestingly, the average of the distribution is roughly equal to one for all categories, showing that on average these categories evolve at the same rate. Therefore, when we average over contexts, there is no significant difference in the speed of evolution of these different parts of the sequence.

To explore the speed of evolution in the different categories, we adopt the same strategy as in Figure 1: we simulate evolutionary trajectories starting from slow, intermediate and fast backgrounds (the same as in Figure 1**E**). We then compute the distance to the starting sequence as a function of time, restricted to the sites that belong to a specific category. Distances are normalized so that a value of one indicates that all amino acids in the considered category are different. Results (Figure 3**E**) confirm that our ranking procedure gives relevant information regarding long-term evolution: the sequence distance to the origin for the context-dependent sites changes at dramatically different speeds if evolution starts from slow, intermediate or fast contexts. On the other hand, not much difference can be seen for the two other categories. Evolutionary speed is therefore strongly affected by epistasis and by the context at a restricted amount of protein sites, while it appears to be well described by a site-independent model at other positions.

### F. Inference of evolutionary distance

An important part of phylogenetic inference consists in inferring the evolutionary distance separating two sequences. For simplicity and because confusion with physical time is not possible in our setting, we refer to parameter *t* as “time” instead of “distance” in the following. Here, we want to understand whether inference done with a model that considers sites as independent becomes biased when epistasis is present in the actual evolutionary model. To do so, we design the following setup. First, we sample an ensemble of *M* = 10 000 sequences {**a**_1_ … **a**_*M*_} from an evolutionary model (profile or Potts) as well as a set of *M* time values {Δ*t*_1_ … Δ*t*_*M*_} from a uniform distribution *U* ([0.01, 2.0]). Then, from each sequence **a**_*i*_, we simulate evolution for time Δ*t*_*i*_ to reach sequence **b**_*i*_. The ensemble {**a**_*i*_, **b**_*i*_, Δ*t*_*i*_} can then be used to assess the accuracy of time inference methods. In our case, we reconstruct the maximum likelihood time Δ*t*_*i*_ from the pair (**a**_*i*_, **b**_*i*_) using the profile model (Material and Methods).

Figure 4**A** shows the reconstructed time as a function of the real time for both profile and Potts as the evolutionary model. On average, the inference with the profile model is very accurate when evolution was also simulated with the profile. However, we note that the time is systematically underestimated when simulating under the Potts model. This underestimation is significant for long times but negligible at short times. This is consistent with previous results, showing that at a given number of substitutions, Hamming divergence from the initial sequence is slower when epistasis is present. Figure 4**B** shows the same data, but this time segregated by the rate of the starting sequence: sequences **a**_*i*_ with an *R*(**a**) in the [0, 1*/*3] quantile form slow contexts, and so on (see Figure 1). For the profile model, this segregation makes no difference, which is not surprising since time inference is performed with the same model as was used to simulate. However, for the Potts model, we observe that for slower contexts the underestimation of time is stronger and significant at smaller times. It is important to stress that the profile model we use for inference has site-specific and sequence-specific rates that are very similar to the Potts, as shown in Figures 1&2. Its equilibrium distribution is also as close as possible to the one of the Potts model, since it has the same site-specific frequencies of amino acids. This systematic bias is therefore an irremediable effect of epistasis.

We try to understand how this bias arises using the ranking of sites according to how context-dependent they are. For each pair of sequences (**a**_*i*_, **b**_*i*_), we again inferred the most likely time with the profile model but restricting the sequences to the *L/*3 most context-dependent, intermediate and least context-dependent sites, that is to the categories shown in Figure 3. In Figure 4**C**, we again plot the inferred time against the real one for each of the three sequence categories, segregating by starting context. It is important to note that the all three categories contain variable sites and each have an average evolutionary rate around one: this guarantees that there are a good amount of substitutions in each category during evolution, making the inference of time possible. Two things can be noted from these plots. First, the underestimation of time is more pronounced for groups of sites that are more context-dependent. In the left-most panel showing the context-independent category, the inferred time is very close to the real one: this indicates that these sites evolve roughly as the profile model predicts, that is independently from each other. On the contrary, deviations are much stronger for the context-dependent category. Secondly, the dependence on the starting context is also stronger when looking at sites that are context-dependent. In the figure, this is materialized by the difference between the three colored curves representing the three starting contexts. This is consistent with Figure 3**E**. These results show that *(i)* inferred evolutionary time is systematically *(ii)* underestimated if epistasis is neglected and underestimation is caused by a set of sites that are strongly context-dependent and is stronger when considering sequences that have a low evolutionary rate.

Finally, we conduct a simple experiment to measure the effect of epistasis on phylogenetic reconstruction using standard tools in the field. We create a set of rooted four-taxon trees with equal branch lengths and diverse heights, defined root-to-leaf distance (Material and Methods). We then simulate evolution on this trees using root sequences of different rates and different evolutionary models: the LG+G4 substitution process as implemented in Alisim [4, 39], and the profile and Potts models. We then infer the phylogeny from the obtained alignments using IQ-TREE [42], using the ModelFinder [43] option to decide what model to use for inference.

Figure 4**D** shows the diameters of the inferred and real trees in the different settings, averaged over replicates of the experiment. Unsurprisingly, inference is on average excellent across the full range of diameters if the LG+G4 model was used as the simulator, as it is then also used for reconstruction. On the other hand, branch lengths tend to be underestimated for large trees in the case of the profile and Potts models, leading to a “saturation” effect in the figure. This effect has already been observed when evolutionary models that model selection are used for simulations while models used for inference ignore it, as in [5] where site-independent models are investigated. This systematic underestimation is therefore not specific to epistasis, but can also be caused by different site-specific frequencies. Where the Potts model strongly differs from the site-independent ones is in the dependence on the starting context. For fast starting contexts, time is surprisingly slightly overestimated for small trees; in contrast, it is strongly underestimated for slow contexts. These results are consistent with the ones of panel **B**, showing that our epistasis indeed leads to biased branch length reconstruction when standard inference tools are used. The four-taxon setup also allows us to test the recovery of topology, with results shown in Supplementary Figure 17. Topology is always correctly recovered when LG+G4 is used to simulate, while errors become frequent for large trees under the profile and Potts model. However, dependence on the context is less clear in this case and it is harder to draw conclusions on the effect of epistasis.

## III. DISCUSSION

We have presented a novel generative protein sequence evolution model based on a Potts model. Its key innovation lies in its continuous-time formulation made possible by the use of the Gillespie [36] algorithm. This allows the model to align with the standards of the phylogenetic literature, while retaining the capacity to produce realistic, functional sequences. With the development of likelihood-free machine-learning based approach to phylogeny, there is a strong need for realistic sequence evolution models [11, 44]. We believe that this work is an important step in this direction.

Our approach relies on several modeling choices. First, the model assumes a single, fixed fitness landscape for an entire protein family, as defined by the Potts model parameters. This assumption implies that the complex web of epistatic interactions that the model captures is static over evolutionary time and across lineages. Changes in the sequence-function mapping – due for instance to a change in cellular environment or of activity – are not considered. A second necessary assumption to make our results biologically significant is that the energy of the Potts model is related to sequence functionality. This has to a large extent proven to be the case: for the families in this paper, Potts models have been used to predict interaction partners or mutational effects [45–47]. However, the model fits statistical patterns in the data that can be unrelated to function: alignment quality, split in different sub-families, and importantly phylogenetic effects. These can give rise to spurious couplings that will influence the dynamics. A caveat to our results is therefore that the energy represents a mixture of different effects: we cannot claim that the epistatic effects that slow down evolution in our model are entirely due to functional constraints. However, successful application of similar dynamical models to experimental data confirms that functional effects play a major role [32, 33]. Among the possible dynamics, the mutation-selection formalism provides the most direct biological interpretation: the energy difference Δ*E*(**a, b**) = *E*(**b**) − *E*(**a**) between two sequences can be interpreted as the product of the effective population size *N*_*e*_ and the selection coefficient of the new variant, connecting to a long tradition of mutation-selection models in phylogenetics [5, 7, 48]. Our results do not qualitatively depend on this choice, as shown in the Supplementary Material. Finally, we incorporate insertions-deletions by treating the gap symbol as an additional amino acid state. This choice has proven effective in the field of generative models, but could be discussed when dealing with evolutionary dynamics as indels and point mutations can have quite different dynamics [49]. This differs from standard phylogenetic techniques which generally treat gaps as missing data [42], and we have argued that it is an improvement in the context of ancestral sequence reconstruction [50].

Our results elucidate both the similarities and stark differences between epistatic and non-epistatic evolutionary models. The qualitative picture is consistent with prior discrete-time epistatic models [29, 31]; continuous time enables quantitative rate analysis and direct connection with phylogenetic inference tools. Surprisingly, we find that the average substitution rates at the sequence and site levels were highly similar between the Potts and profile models. However, these average rates are only relevant for very long evolutionary trajectories where the context is averaged over. The effects of epistasis become profoundly apparent when examining specific sequence contexts, where we find site-specific substitution rates to strongly differ between the Potts and profile models. Another observation is that the speed of evolution is highly dependent on the starting sequence, with low-energy sequences *(i)* accumulating fewer substitutions per unit time and *(ii)* needing a significantly higher number of substitutions to diverge from the initial sequence. We further demonstrated that this epistatic influence is not uniformly distributed across the protein sequence. Indeed, some positions are highly constrained by the rest of the sequence – with evolutionary rates ranging from vanishing to high depending on the context – while others evolve in a largely independent manner. Intriguingly, the long-term average evolutionary rates for these two groups of sites are relatively close, masking the critical role of epistasis when only average quantities are considered. This uncovers interesting features of evolution in a complex landscape while showing that site-independent models are unable to capture them.

A central finding with important implications for phylogenetic inference is that ignoring epistasis leads to a systematic underestimation of evolutionary time. This bias is not uniform; it is significantly stronger *(i)* for slow-evolving sequences and *(ii)* when inference is based on highly context-dependent sites. Importantly, this bias arises even if the site-independent model that is used for reconstruction has correct average site-specific rates. Even if we do not explore it here, it is reasonable to think that these biases will impact full-scale phylogenetic reconstruction. For instance, epistasis could cause a uniform compression of all branch lengths or distort the tree topology by preferentially shortening branches leading to specific clades. Clades that have evolved in a context of high functional constraint (our “slow” sequences) might be disproportionately affected, making recent evolutionary radiations appear even more recent than they are and potentially obscuring their true chronological order. As it is now well established that epistasis is ubiquitous in protein evolution, understanding how it affects phylogenetics and how we can correct for it becomes an important research direction to which we hope this work will contribute.

## IV. MATERIAL AND METHODS

### A. Evolutionary model

In general, the evolutionary model gives the rate of transition between any two sequences **a** and **b**. We assume that this rate is non-zero only between sequences separated by one amino acid mutation, and define *N*(**a**) as the set of sequences that are one mutation away from **a**. Consequently, a given row *Q*_**a**:_ of the transition rate matrix has only (*L* − 1)*q* non-zero values where *q* = 21 is the number of possible amino acids plus the gap symbol. The full model is therefore specified by

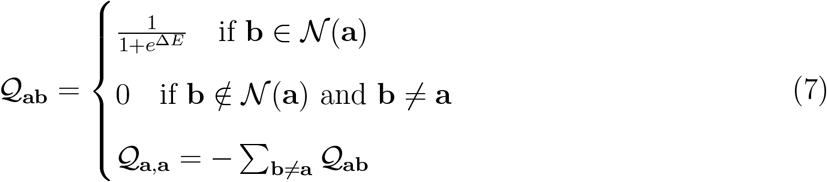

where Δ*E* = *E*(**b**) − *E*(**a**), and where the expression for *Q*_**a**,**a**_ ensures that rows of *Q* sum to 0 as should be the case for a continuous time Markov chain.

We use a normalization constant Ω to set the timescale of the process. Following common practice in modeling sequence evolution, we want the average rate of substitutions per position to be one, that is

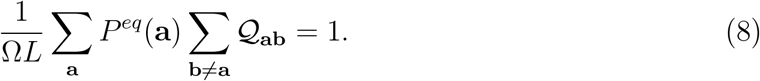

This is achieved by setting Ω to

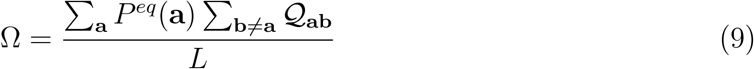

This average over sequences **a** can be calculated either using a sample from *P*^*eq*^ obtained by standard MCMC, or using natural sequences to compute the average. The matrix *Q* is then scaled by a factor Ω^*−*1^, so that transition probabilities take are (informally) of the form Exp 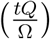. Ω therefore represents the amount of “physical” time corresponding to one unit of evolutionary time. Supplementary Figure 18 shows the value of Ω for family PF00072 and different types of Monte Carlo dynamics. We see that for each dynamic, the value of Ω is higher for the Potts model than for the profile, indicating that epistasis also has the effect of changing the scaling from evolutionary time to “physical” time. In the main text and in the rest of this section, this scaling is implicit and we lighten notation by omitting Ω.

### B. Protein families

This work uses three protein families: the response regulator (PF00072), the PDZ domain (PF00595) and the beta-lactamase (PF13354).

- The alignments for the PF00072 and PF00595 family are obtained in the same way as in [51]: they are downloaded from Pfam [37] and filtered for sequences with more than six consecutive gaps. As the PF00072 alignment is very large (more than 800 000 sequences), we sub-sample it by randomly picking one sequence out of ten.
- The alignment for the PF13354 family is the same as the one used in [32]. It was obtained by running the `hmmsearch` command of HMMER [52] against Uniprot [53], and then filtering out sequences with 20 gaps or more or that have duplicates.

### C. Inferring Potts models

There are a variety of ways to infer the parameters {*J, h*} from a multiple sequence alignment. Methods such as pseudo-likelihood maximization (plmDCA) or mean-field (mfDCA) are fast and accurate enough to perform structural contact or mutational effect prediction [19, 20, 46]. However, the resulting generative model is not accurate in the sense that the statistics of the model in Eq. 1 do not precisely reproduce the statistics of the natural sequences [21].

We therefore infer the Potts models in this work using Boltzmann-Machine algorithm (bmDCA), which is computationally more expensive [21]. For all families, we used the adabmDCA implementation of [38] with *l*_2_ regularization set to 0.01. The implementation partially corrects the statistics of the natural sequences for phylogenetic biases by applying a reweighting technique originally proposed in [19]. For PF00072 and PF00595 models, default parameters were used. PF13354 has longer sequences, making training more difficult, and we adapted the duration and number of Monte Carlo chains gradually during the training until convergence of the algorithm.

On the other hand, profile models were inferred by simply using the analytical formula

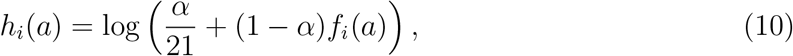

where *f*_*i*_(*a*) is the frequency at which amino acid *a* is found in column *i* of the alignment. Parameter *α* is a pseudocount serving for regularization, and is set to 0.01.

### D. Simulating with Gillespie algorithm

We use Gillespie’s algorithm to sample from our model and simulate evolution [36]. Given a starting sequence **a**, this is done in three steps:

1. compute the total rate of substitution away from **a**: *R*(**a**) = ∑ _**b***∈N* (**a**)_ *Q*_**ab**_;
2. sample the time *t* to the next substitution from an exponential distribution of parameter *τ* = 1*/R*(**a**);
3. choose the next substitution: position *i* and new amino acid *b* are chosen with probability

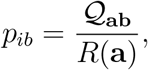

where **b** is equal to **a** at all positions but *i*, where it is equal to *b*.

From Eq. 8 we get that the average rate *R*(**a**) over starting sequences is *L*, that is a rate 1 per sequence position on average.

### E. Ranking of positions based on context-dependent rates

Our aim is to identify sets of positions that *(i)* have an evolutionary rate that strongly depends on the context and *(ii)* have a high or low in similar contexts. In order to do this, we note *A*_*K*_ an arbitrary set of *K* positions and define for any sequence **a** the partial sum of context rates as

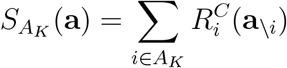

where 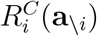 is defined in Eq. 6. For each *K*, we now want to find the set of positions *A*_*K*_ that maximizes the variance of 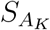 across natural sequences. Note that this variance depends on both the variance and covariance of the context rate:

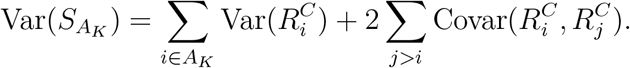

We estimate the best set *A*_*K*_ in a greedy way: starting from the set *A*_*K−*1_, we add to it the position that marginally maximizes the variance. The order in which we add positions is the ranking discussed in Figure 3. To evaluate how accurate the greedy approach is, we also try to optimize the set *A*_*K*_ using a more presumably precise simulated annealing approach. Results using the two methods are close to identical.

### F. Inferring time with the profile model

We show in the Supplementary Section A 2 that when using a profile model, the transition probability between two sequences in time Δ*t* can be written as

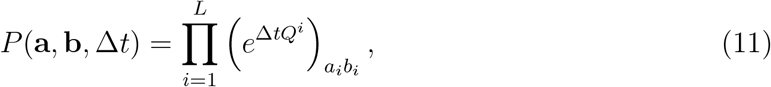

where the *Q*^*i*^ are 21 × 21 giving the transition rates between amino acids at position *i*, and 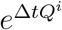 is the matrix exponential. Given two sequences **a** and **b**, we infer the evolutionary time between them by maximizing the log-likelihood

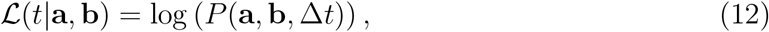

which is simply a sum over *L* terms that each involve the states *a*_*i*_ and *b*_*i*_. To speed-up the optimization, we preemptively diagonalize the matrices *Q*^*i*^ that depend only on the model in order to quickly compute their exponential. We perform the optimization with the LBFGS algorithm along with automatic differentiation [54, 55].

### G. Inference for four-taxon trees

We measure the performance of phylogenetic inference software when data was generated under the Potts model using a simple four-taxon experiment. Given an evolutionary model, we simulate data on a four-taxon tree of the form `((A,B),(C,D))` where A, B, C and D are the leaves. All branches have the same length so that the tree is simply parametrized by its height *T* (root to leaf distance). We pick three sequences of the PF00072 family to place at the root to start evolution from, picked from quantiles (10%, 50%, 90%) of the distribution of *R*, respectively defining slow, intermediate and fast contexts. Evolution is simulated using Alisim [39] with the *LG* + *G*4 model with shape parameter 1 for the Gamma distribution, the profile model and the Potts model. For each model, root sequence and tree diameter, we perform 25 simulations and obtain as output as many alignments of leaf sequences.

Phylogenetic inference is performed using IQ-TREE [42] using default parameters and the model finder [43] to select the best model for reconstruction. Before inference, we remove all alignment positions that contain at least one gap, and this to avoid two potential problems due to gaps. First, the Potts and profile models simply treat gaps as a 21st state, which is inaccurate for insertion-deletion dynamics. Secondly, it is common in phylogenetic software to treat gaps as “missing data”, which is also inaccurate. The inferred diameter of the tree is computed by averaging the leaf to leaf distances *d*_*AC*_, *d*_*AD*_, *d*_*BC*_, *d*_*BD*_. Only trees with a correct topological inference are used in the results displayed in Figure 4.

## CODE AVAILABILITY

The code used to simulate our evolutionary model is published as a Julia package at https://github.com/PierreBarrat/PottsEvolver.jl, along with a documentation. The scripts used to generate evolutionary trajectories and the figures of this paper are available at https://github.com/PierreBarrat/ContinuousPottsData. The simulated evolutionary trajectories can be found at https://doi.org/10.5281/zenodo.18221837.

## ACKNOWLEDGMENTS

AP acknowledges financial support from “Explainable Models for Protein Design”, funded by the MIUR Progetti di Ricerca di Rilevante Interesse Nazionale (PRIN) Bando 2022 - grant 2022TE5B7X, “Centro Nazionale di Ricerca in High-Performance Computing, Big Data and Quantum Computing” (ICSC), and support from the European REA, Marie Sk-lodowska-Curie Actions, grant agreement no. 101131463 (SIMBAD). AP and PBC acknowledge “FAIR - Future Artificial Intelligence Research” project, and received funding from the European Union NextGenerationEU (Piano Nazionale di Ripresa e Resilienza (PNRR)–Missione 4 Componente 2, Investimento Grants No. 1.3–D.D. 1555 11/10/2022, and No. PE00000013). This paper reflects only the authors’ views and opinions, neither the European Union nor the European Commission can be considered responsible for them.

## Supplementary Material

## Appendix A: Supplementary methods

### 1. Different dynamics

In the main text, we use our model with Glauber dynamics (Eq. 2). Other choices are possible: any transition rate matrix Q that satisfies the detailed balance conditions will be compatible with the generative distribution *P*^*eq*^. In the supplementary figures below, we show results for three alternative dynamics (defined here in the case **b** ∈ *N*(**a**)):

- Metropolis dynamics:

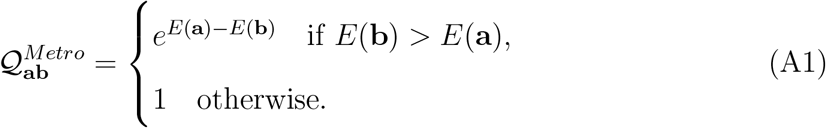
- Square-root rule:

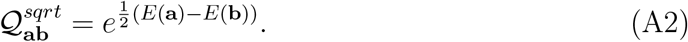

This rule has been used in the past in the context of evolutionary models [26]. We do not know of a proper name for it, and choose to call it this way because the transition rate is equal to the square-root of the ratio *P*^*eq*^(**b**)*/P*^*eq*^(**a**).
- Gibbs dynamics:

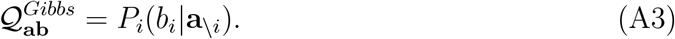

The rate of transition to state *b*_*i*_ at position *i* is equal to the conditional probability of *b*_*i*_ given the rest of the sequence. Note that the Gibbs rule is the only one that does not depend on the initial state *a*_*i*_ at position *i*.
- Mutation-selection: this dynamic has been proposed in the context of phylogenetic inference to model site-specific selective effects [5, 7]. The rates are given

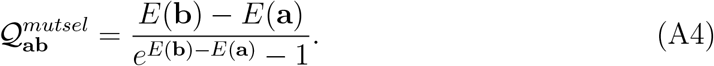

It is of particular interest as it connects exponential distributions such as the Potts model (Eq. 1) to population genetics theory [48]. Indeed, if we assume the Potts energy difference *E*(**b**) − *E*(**a**) to be proportional to the product of the effective population size *N*_*e*_ and minus the selective advantage of sequence **b** with respect to a background population carrying sequence **a**, then under certain conditions the fixation probability of the newly appeared mutant **b** is

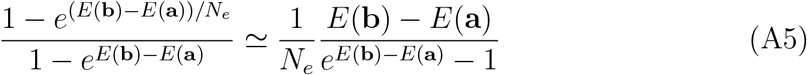

according to the expression derived by Kimura [56]. The approximation in the r.h.s is valid for small selective advantages, that is *E/N*_*e*_ ≪ 1. While the effective population size is unknown, it is effectively absorbed into time as we rescale the rates Eq. 9.

Note that in the work originally introducing this dynamic [5], the rates 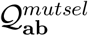 are multiplied by a mutation propensity between amino-acids *µ*_*ab*_. Such mutational biases could straightforwardly be integrated in our framework, as could be constraints linked to the genetic code.

### 2. Profile model

We want to show that the transition rate matrix of Eq. 7 is compatible with non-epistatic evolution. To do so, we design a profile evolutionary model where each sequence position *i* follows its own continuous time Markov chain with rate *µ*_*i*_ and a *q* × *q* transition rate matrix *Q*^*i*^, and show that its sequence to sequence transition rate matrix is equal to the *Q* of the main text. The transition probability between two sequences **a** and **b** in time *t* is

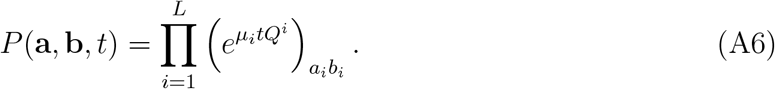

Each matrix *Q*^*i*^ implicitely defines a stationary probability *π*_*i*_(*a*) for amino-acids at site *i*, such that for any *b*,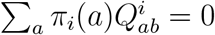. We assume that the *Q*^*i*^ and *µ*_*i*_ are normalized:

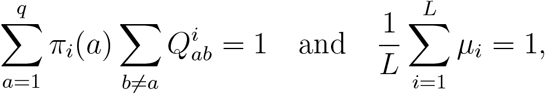

so that position *i* evolves on average at rate *µ*_*i*_ and the average rate at each position is one.

The corresponding *sequence to sequence* transition rate matrix *Q* is defined by

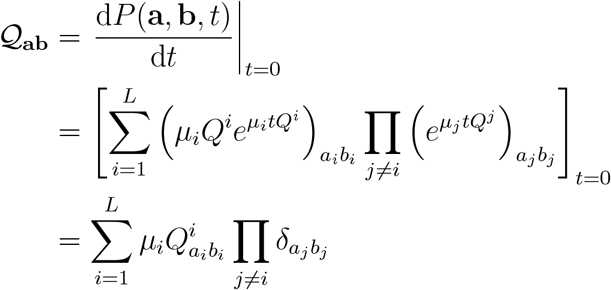

with *δ*_*xy*_ being the Kronecker delta. Therefore the transition rate vanishes for sequences that are more than one mutation away and we can write

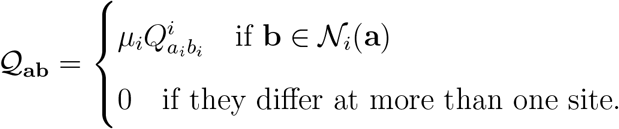

For this to match the matrix of the main text in Eq. 7, we need that for any sequences **a** and **b** ∈ *N*_*i*_(**a**), the transition rate *Q*_**ab**_ depends only on the states *a*_*i*_ and *b*_*i*_ at position *i*.

This is necessary as on the right hand side of the above equation, 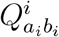 is independent of the context. Note that for the Glauber dynamics, but also for other dynamics described below, *Q* automatically has this property if the equilibrium distribution *P*^*eq*^ is factorized, which is the case when we use a profile model.

More precisely, models in Eq. 7 and Eq. A6 are the same if we have

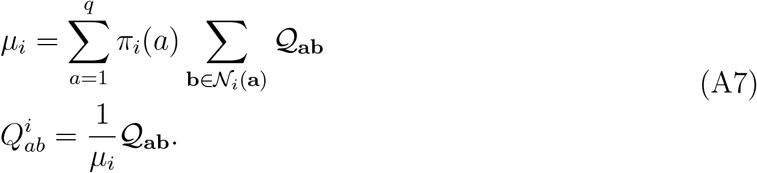

where **a** is any sequence having state *a* at position *i*, and **b** is equal to **a** except at position *I* where it is in state *b*.

What we have shown here is that applying the model of the main text to a factorized distribution *P*^*eq*^ results in a factorized dynamics (Eq. A6) with inhomogeneous rates *µ*_*i*_.

### 3. Link between substitution rate and conservation

In Figure 2, we saw that the average substitution rates of sites strongly is strongly correlated to conservation. This can be explained analytically in the case of the profile model.

In this case, if we call *π*(*a*) the probability of a given site to be in state *a*, then its average rate in the profile model will be

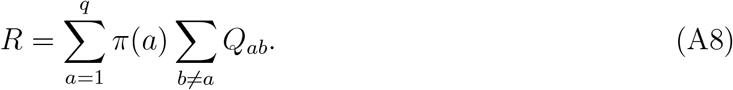

where *Q* is the site-specific *q* × *q* transition rate matrix as in A6 and *q* = 21 is the number of states. The result of the calculation will of course depend on the type of dynamics used. The simplest formula turns out to be obtained in the case of Gibbs, where *Q*_*ab*_ = *π*(*b*). We then obtain

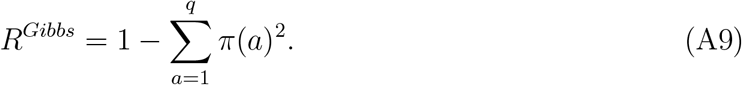

The expression 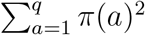 is a measure of how “concentrated” the probability distribution *π* is: it evaluated to one if one state gets all the mass, and to 1*/q* if all states are represented equally. It is therefore no surprise that *R*^*Gibbs*^ strongly correlated with the entropy *S* = − _*a*_ *π*(*a*) log *π*(*a*).

In fact, a similar calculation can be done for all dynamics, leading to the following results:

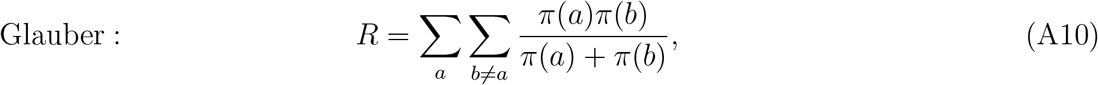

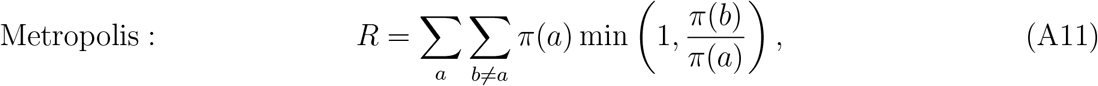

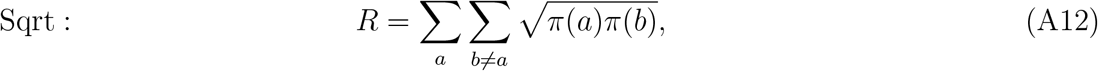

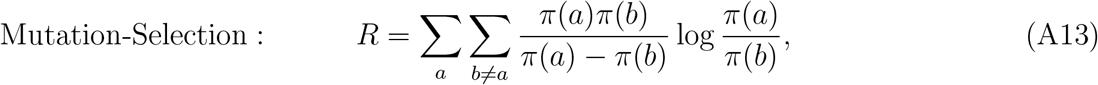

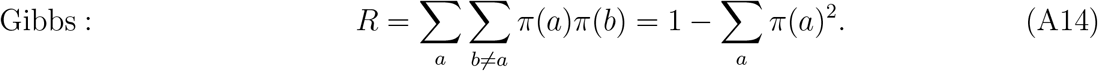

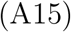

While some of these expressions are more complicated, they are all strongly correlated to the entropy as can be observed in Supplementary Figure 21.

### 4. Distribution of context-dependent site rates

In Figure 2 of the main text, we discuss distribution of average site rates ⟨*R*_*i*_⟩. When fitting a Gamma distribution to this distribution, we find a shape parameter *α* = 2.2 in the case of PF00072. This is a relatively high value, indicating a low variance of rates: in comparison, [4] found values of *α* typically around 1 for Pfam alignments [37]. In this section, we explore in more details the links between average and context-dependent rates and the shape parameter *α* by means of the variance decomposition.

In a deep phylogeny where the context changes enough along the branch lengths, the observed rate at site *i* should indeed be ⟨*R*_*i*_⟩, where we remind that the average is taken over natural sequences. At the other extreme, in a shallow tree, we may consider the context to be fixed so that the rate at site *i* will be the context-dependent rate 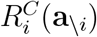 of Eq. 6. First, we note that the average of 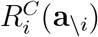 over contexts is exactly equal to the average rate:

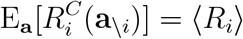

where E_*x*_ represents the average over quantity *x*. The shape parameter of the Gamma fit of Figure 2 will therefore be related to the variance 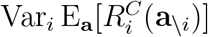. On the other hand, in a shallow phylogeny with an MRCA having sequence **a**, the observed variance in site-specific rates will be 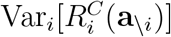. The average variance, averaged over contexts, will therefore be 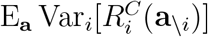. These two quantities can be related using the variance decomposition formula:

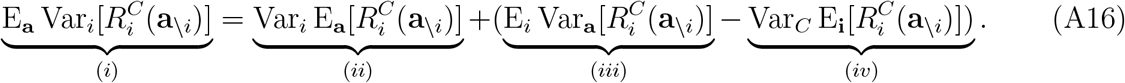

Here is the interpretation of these different terms.

i. Expected variance of context-dependent site-specific rates, over contexts. This is arguably what would be observed in a shallow phylogeny.
ii. Variance over sites of the context-averaged site-specific rates.
iii. Average context-dependence of sites: the term 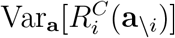 quantifies how variable the rate of site *i* is accross contexts. It corresponds to the average over sites of the variance of histograms in Figure 3**B**.
iv. Variance over contexts of average site rates. Note that with our definition, the average site rate 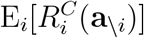is not strictly equal to the scaled sequence rate *R*(**a**) of Eq. 4. However, the two quantities are very close in practice. This term is on the same order of magnitude as the variance of the histogram in Figure 1**A**.

Because of the two terms in parenthesis, we expect different values of the shape parameter *α* when looking at average or context-dependent rates. Visual inspection of Figure 1**A** and Figure 3 of the main text suggests that 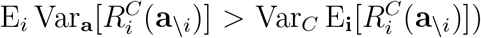, so that the expected variance in site rates in a given context is higher than the variance of context-averaged rates. This is verified quantitatively, since the four terms are respectively equal to 0.67, 0.43, 0.3 and 0.06 in the case of PF00072. Figure 12 shows how these results translate in the more familiar terms of Gamma distribution and shape parameters. In panel **A** it shows the distribution of rates in three specific contexts, with values of *α* that are more typical. This is confirmed exhaustively in panel **B** which shows the distribution of *α* over 1 000 natural contexts.

It is important to note that this discussion is completely irrelevant for the profile model. In this case, since rates do not depend on the context, terms (*iii*) and (*iv*) vanish and the different ways to measure site-heterogeneity coincide. In [5], it is argued that heterogeneous site rates are caused by the fitness landscape, and a non epistatic model is proposed for the landscape. We show here on a limited set of protein families that this gives rise to values of *α* that are above what is typically observed.

## Appendix B: Supplementary figures

**Figure S1.**
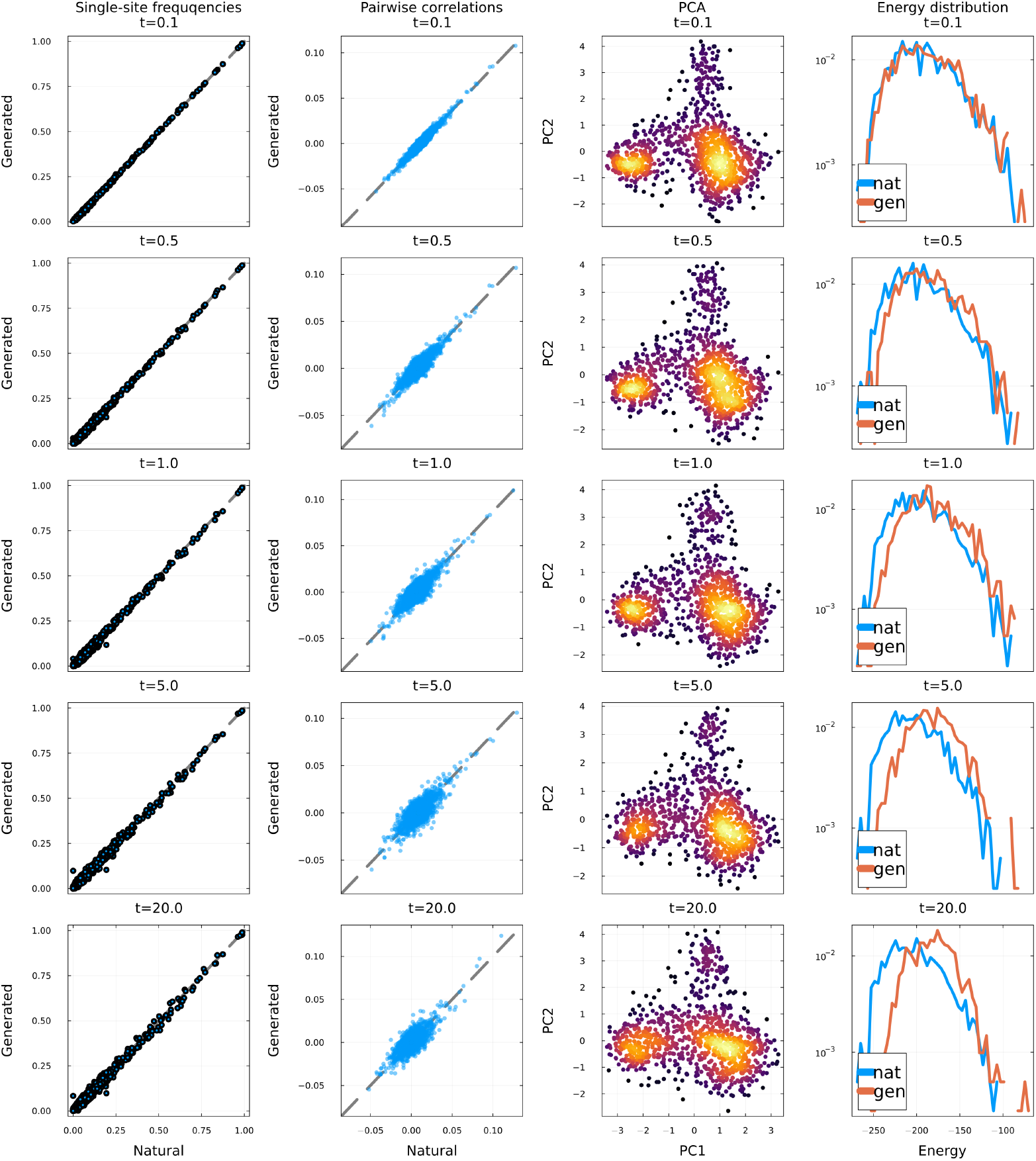
Estimation of the generative capacities of our model. *M* = 1000 evolutionary trajectories are started from different natural sequences of the PF00072 family, and sampled at five different times (0.1, 0.5, 1.0, 5.0, 20). At each time, statistical properties of the *M* evolved sequences are evaluated. **First column**: Column-wise amino acid frequencies of the natural (*x*-axis) and generated sequences (*y*-axis). **Second column**: Pairwise connected correlation between MSA columns of the natural (*x*-axis) and generated sequences (*y*-axis). **Third column**: Projection of generated sequences onto the first two principal components of the MSA of the natural sequences. Points are colored by density. **Fourth column**: Distribution of the Potts statistical energy (Eq. 1) of the natural and generated sequences. *y*-axis uses a logarithmic scale. The slight shift in histograms is caused by the regularization of the model parameters during training [12].

**Figure S2.**
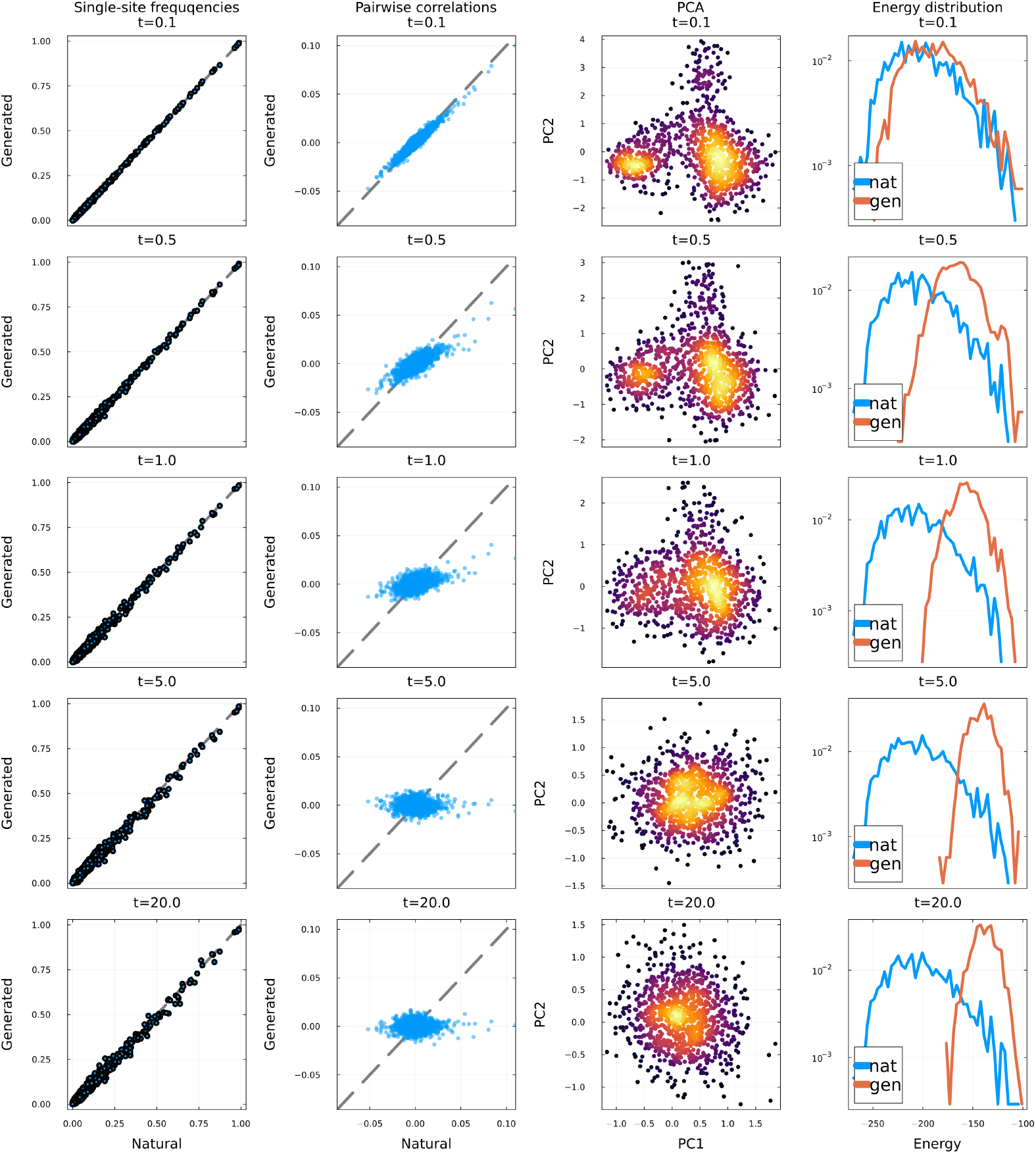
Estimation of the generative capacities of the profile model. *M* = 1000 evolutionary trajectories are started from different natural sequences of the PF00072 family, and sampled at five different times (0.1, 0.5, 1.0, 5.0, 20). At each time, statistical properties of the *M* evolved sequences are evaluated. **First column**: Column-wise amino acid frequencies of the natural (*x*-axis) and generated sequences (*y*-axis). **Second column**: Pairwise connected correlation between MSA columns of the natural (*x*-axis) and generated sequences (*y*-axis). **Third column**: Projection of generated sequences onto the first two principal components of the MSA of the natural sequences. Points are colored by density. **Fourth column**: Distribution of the Potts statistical energy (Eq. 1) of the natural and profile-generated sequences. *y*-axis uses a logarithmic scale.

**Figure S3.**
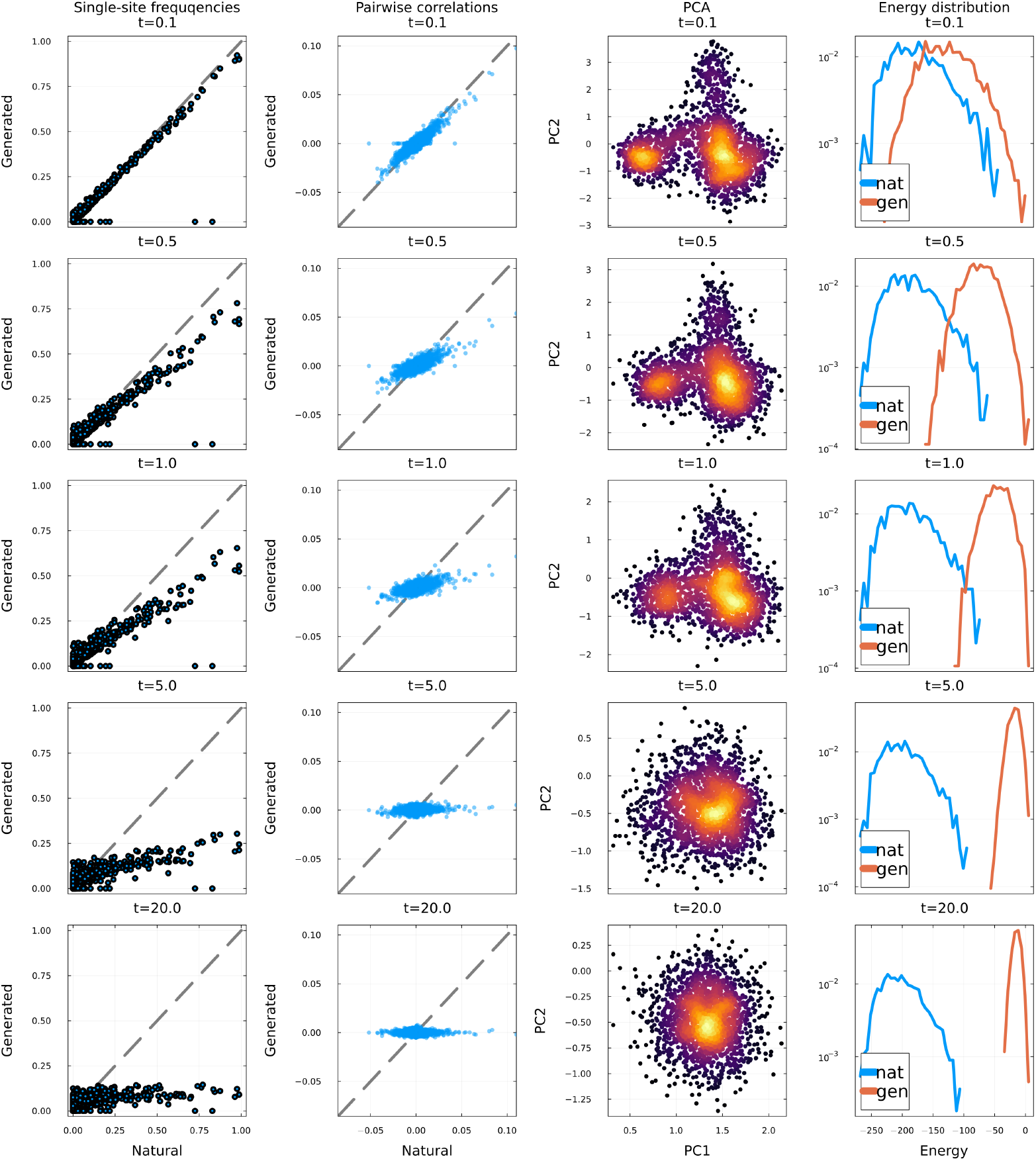
Estimation of the generative capacities of LG+G4 model. *M* = 1000 evolutionary trajectories are started from different natural sequences of the PF00072 family, and sampled at five different times (0.1, 0.5, 1.0, 5.0, 20). At each time, statistical properties of the *M* evolved sequences are evaluated. **First column**: Column-wise amino acid frequencies of the natural (*x*-axis) and generated sequences (*y*-axis). **Second column**: Pairwise connected correlation between MSA columns of the natural (*x*-axis) and generated sequences (*y*-axis). **Third column**: Projection of generated sequences onto the first two principal components of the MSA of the natural sequences. Points are colored by density. **Fourth column**: Distribution of the Potts statistical energy (Eq. 1) of the natural and LG+G4 generated sequences. *y*-axis uses a logarithmic scale.

**Figure S4.**
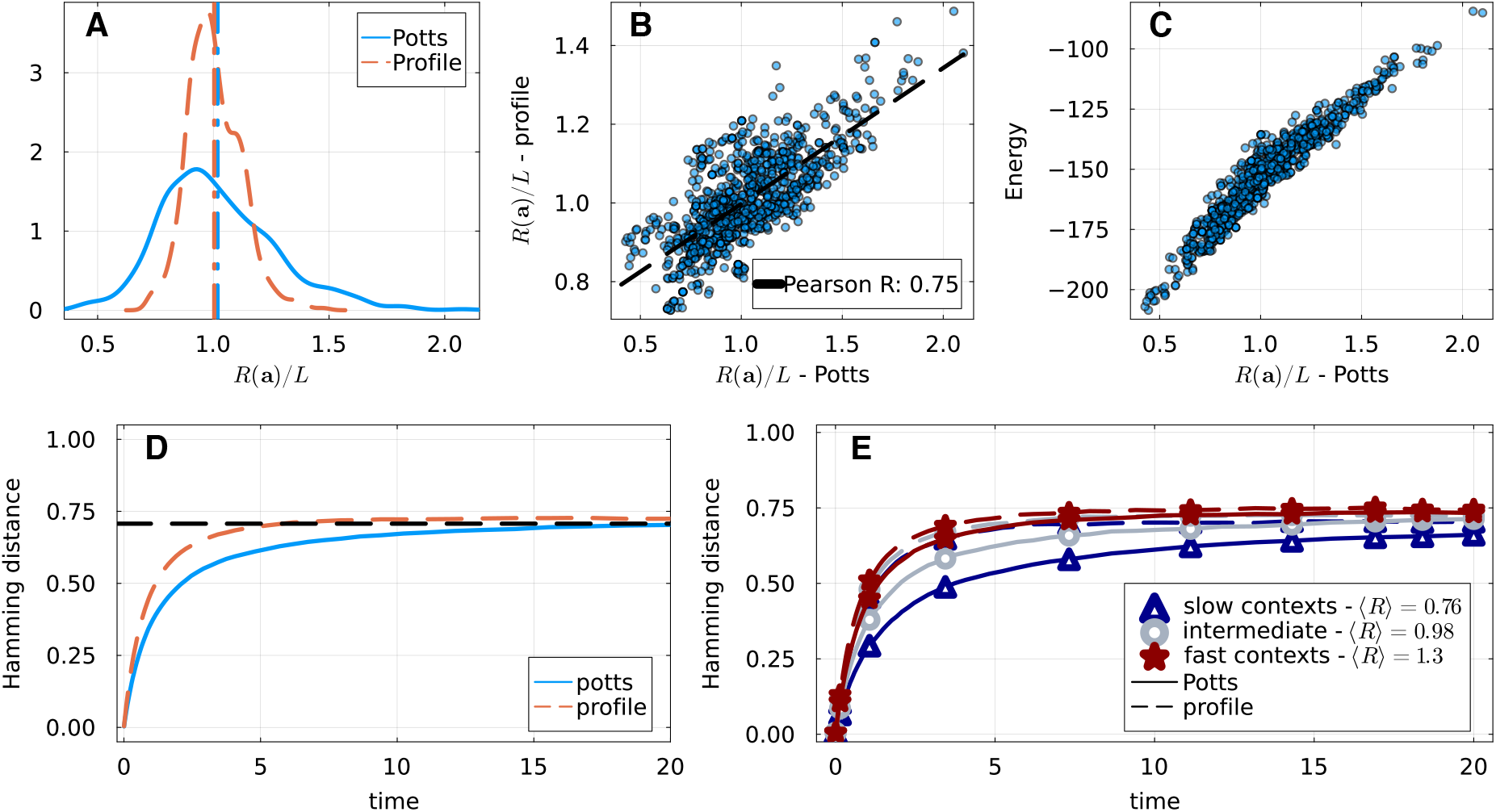
Equivalent to Figure 1 of the main text for the PF00595 family.

**Figure S5.**
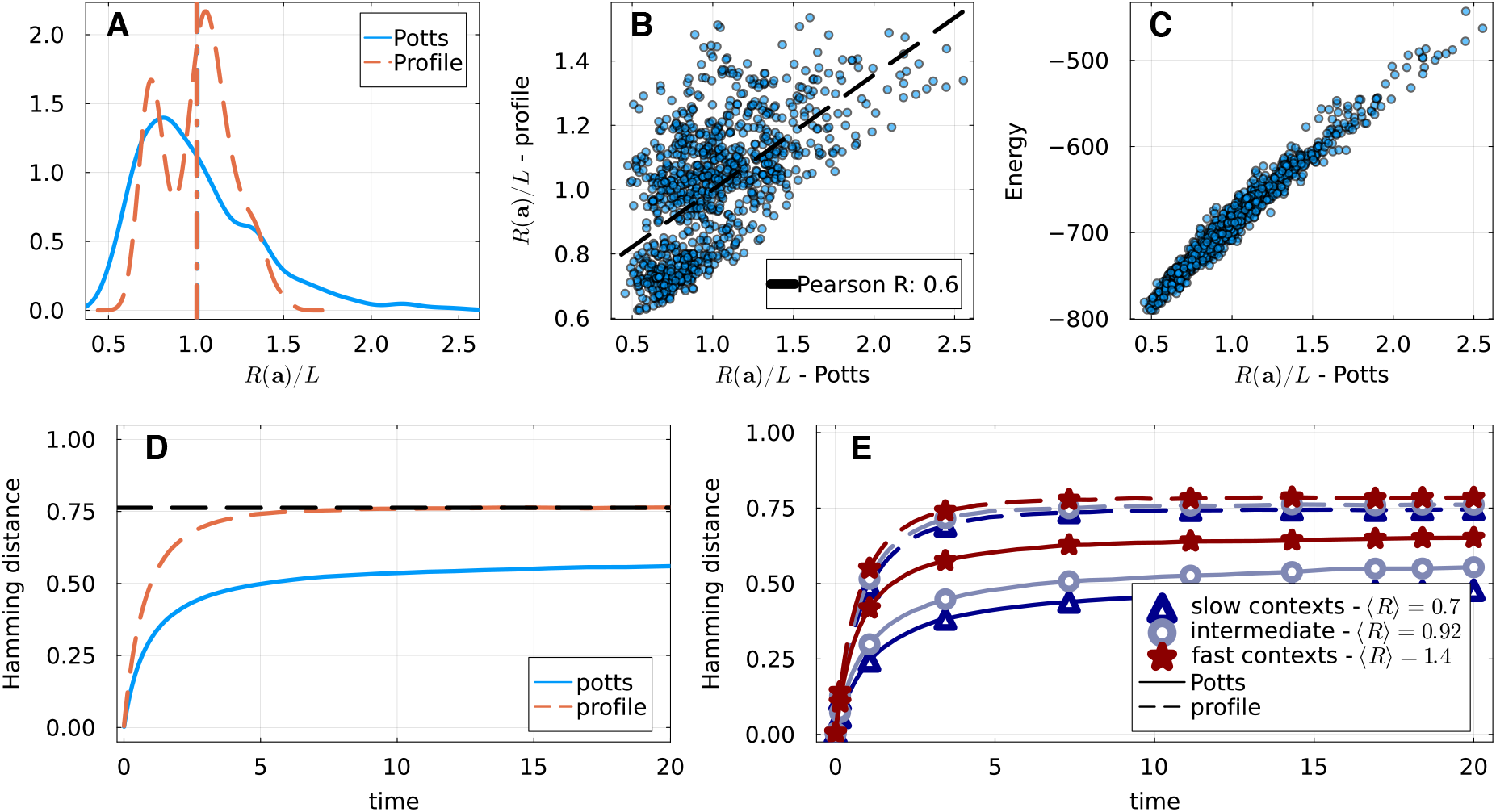
Equivalent to Figure 1 of the main text for the PF13354 family.

**Figure S6.**
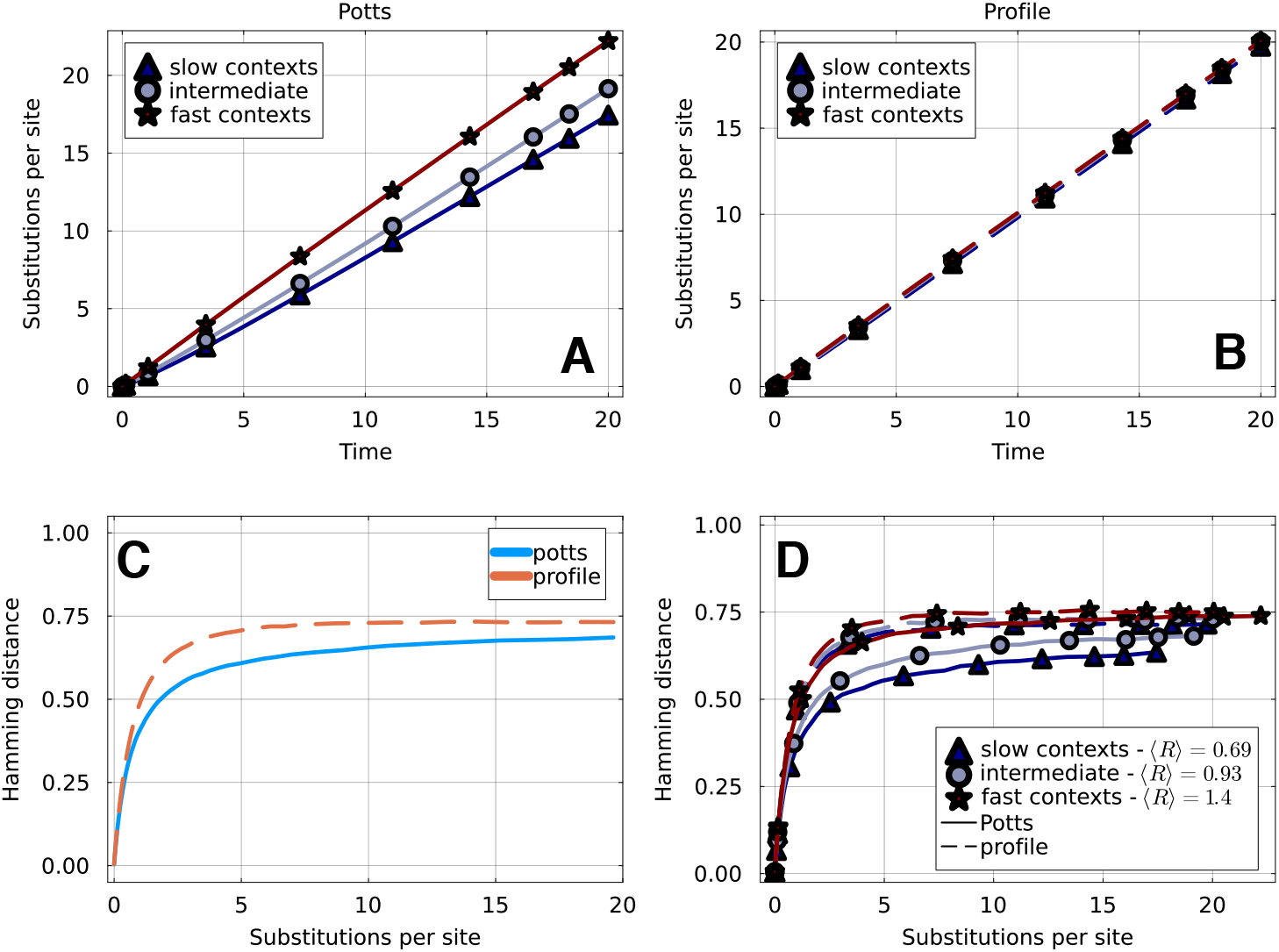
Panels **A&B**: number of substitutions as a function of evolutionary time, for the Potts and profile models. Consistently with Figure 1**A**, we see that rates are more disperse across contexts in the case of the Potts model. **C** Average sequence divergence versus number of substitutions for evolution simulated with a Potts and profile models. The average is over a set of 100 starting sequences and 5 realizations for each starting sequence. Sequence divergence is computed relative to the initial sequence. Equivalent to Figure 1**D** with substitutions instead of time. **D** Average sequence divergence versus substitutions segregated by the category of the starting sequence: slow ([0, 0.33] quantile), intermediate ([0.34, 0.66] quantile) and fast ([0.67, 1.] quantile). The scaled average rate R(a) of initial sequences in each category is indicated in the legend.E quivalent to Figure 1**E** with substitutions instead of time.

**Figure S7.**
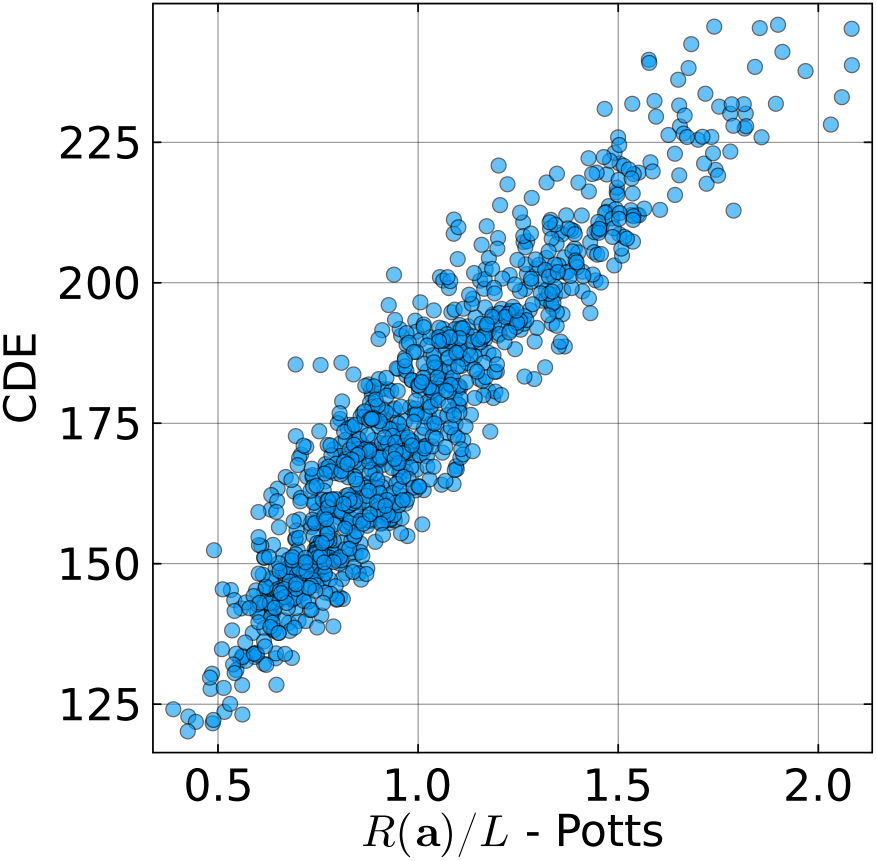
Context dependent entropy (CDE) versus sequence specific rate of evolution in a Potts model for members of the PF00072 family. We define the CDE of a sequence as the sum of the context dependence entropies at each position *i* in the sequence, themselves defined as in [31]. Context dependent entropy has been shown to be good proxy for the velocity of evolution at each position of the sequence.

**Figure S8.**
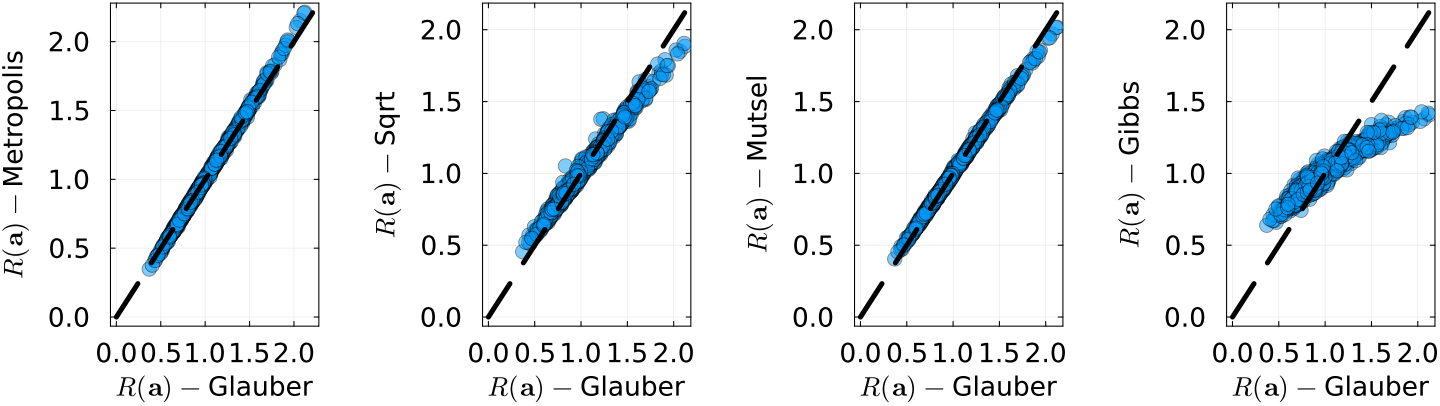
Sequence evolutionary rate *R*(**a**) for different dynamics (scaled by the sequence length). The horizontal axis shows Glauber dynamics used in the main text, and the vertical axes respectively show Metropolis, square-root, mutation-selection and Gibbs dynamics. Rates are very correlated in every case.

**Figure S9.**
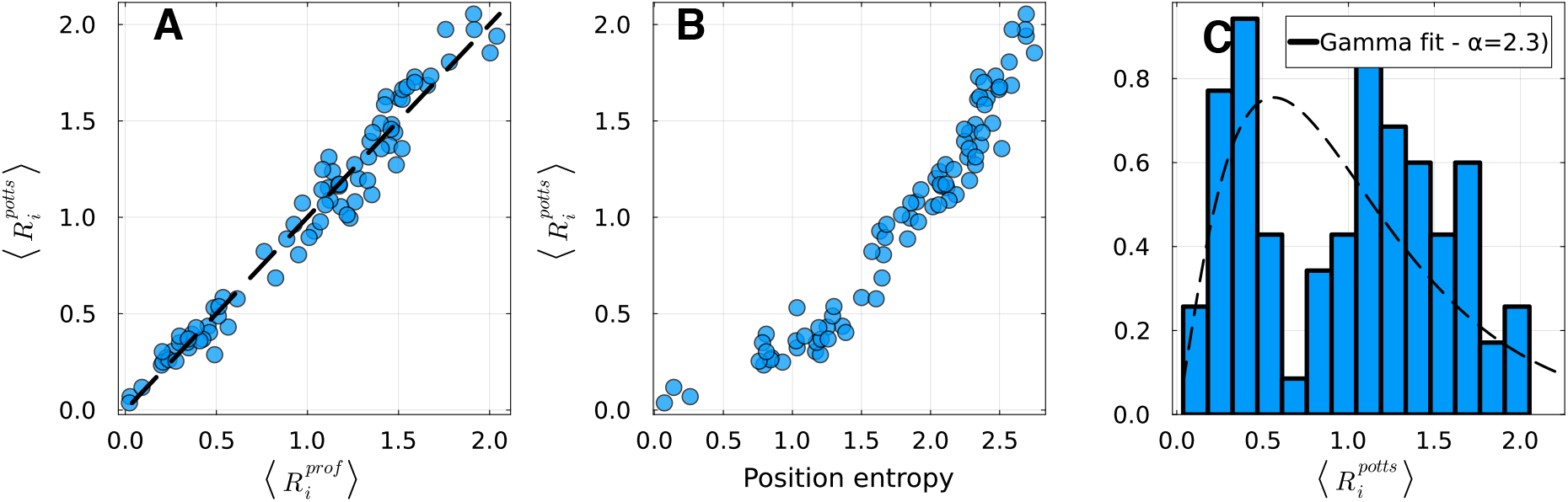
Equivalent to Figure 2 of the main text for the PF00595 family.

**Figure S10.**
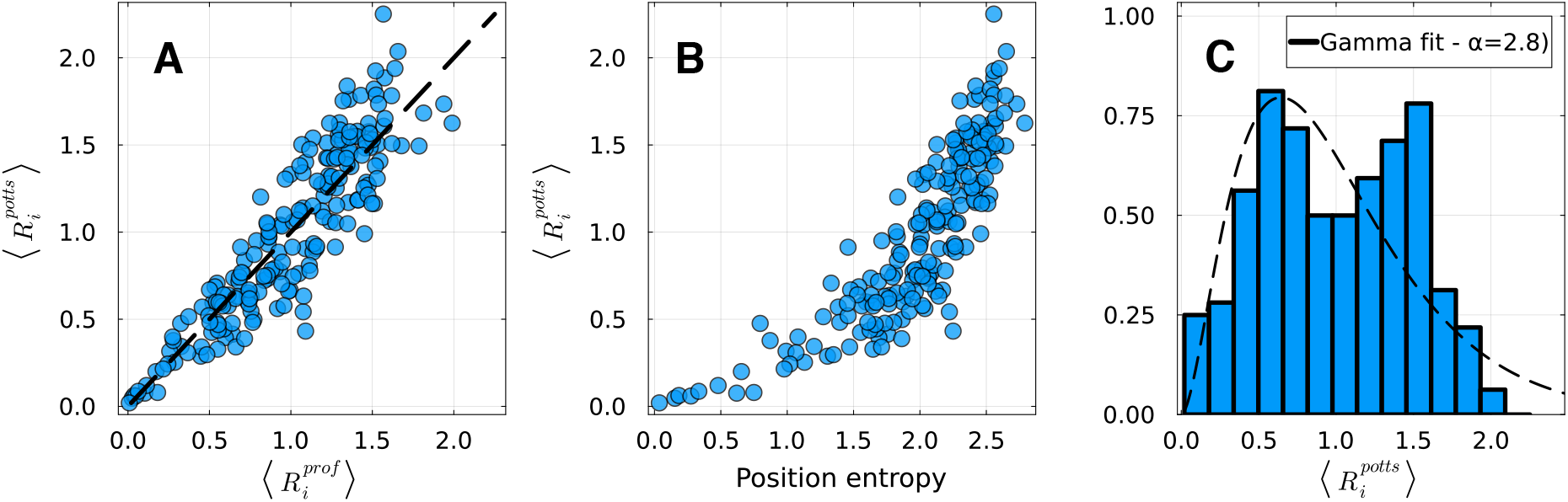
Equivalent to Figure 2 of the main text for the PF13354 family.

**Figure S11.**
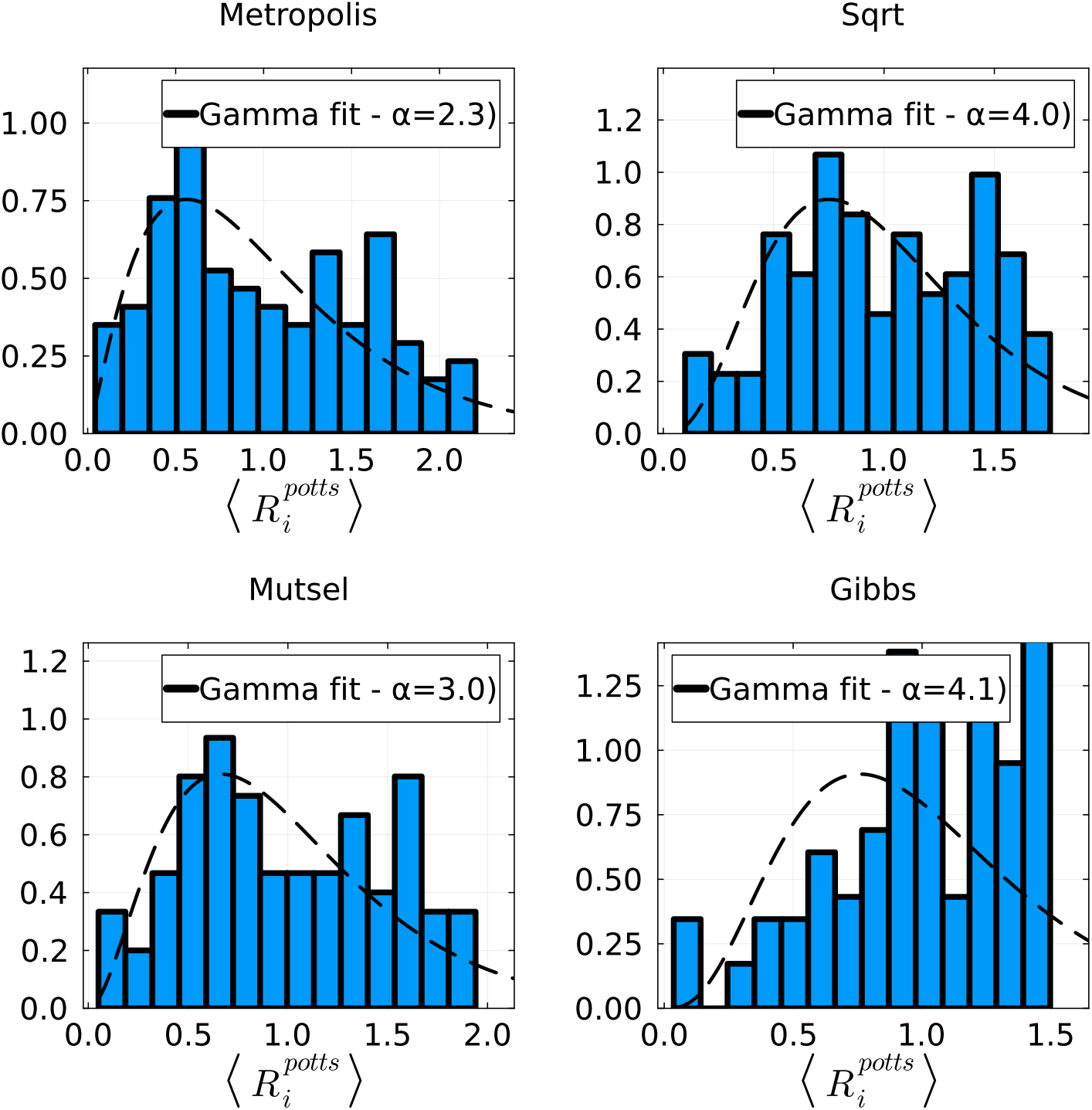
Distribution of average site-specific rates for family PF00072 and different dynamics. This is equivalent to the Figure 2**C** of the main text. The dashed line indicates the maximum likelihood fit with a Gamma distribution.

**Figure S12.**
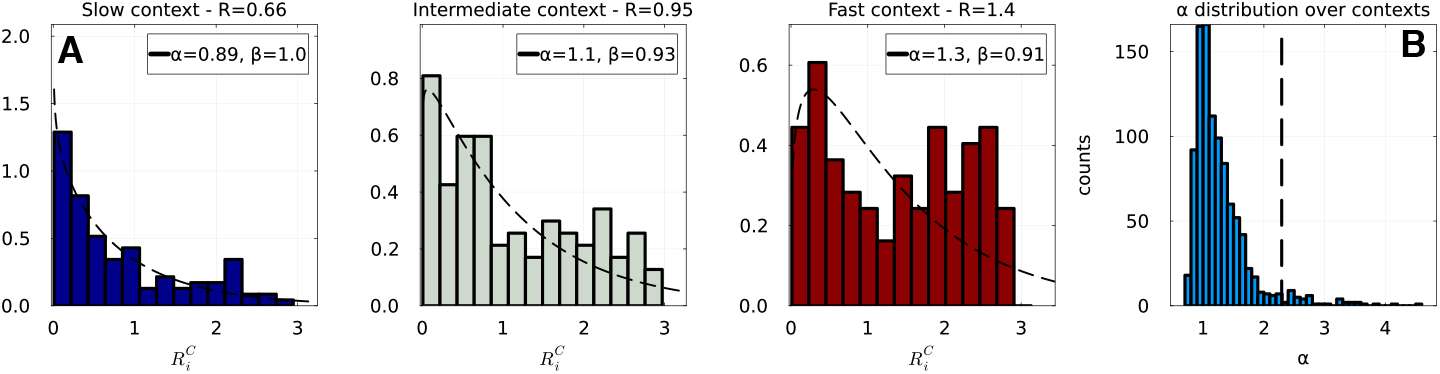
**A**: Distribution of context-dependent site-specific rates 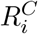 for three different contexts corresponding to quantiles [0.1, 0.5, 0.9] of the sequence rate distribution (PF00072). For each context, we fit a Gamma distribution to the histogram and show the shape and scale parameters *α* and *θ* in the label. This illustrates the fact that rates in a given context give rise to a smaller shape parameter *α* with respect to Figure 2**C**. Note that the averages of these distributions are not one, as some contexts are slower than others in the sense that they accumulate less substitutions per unit time. However, this only affects the scale parameter *θ* and not *α*. **B**: Distribution of inferred shape parameters *α* for contexts coming from 1 000 randomly selected natural sequences (PF00072). The vertical bar indiciates the value found in Figure 2**C** which corresponds to the profile model, or to the Potts model if one averages site-specific rates over contexts first.

**Figure S13.**
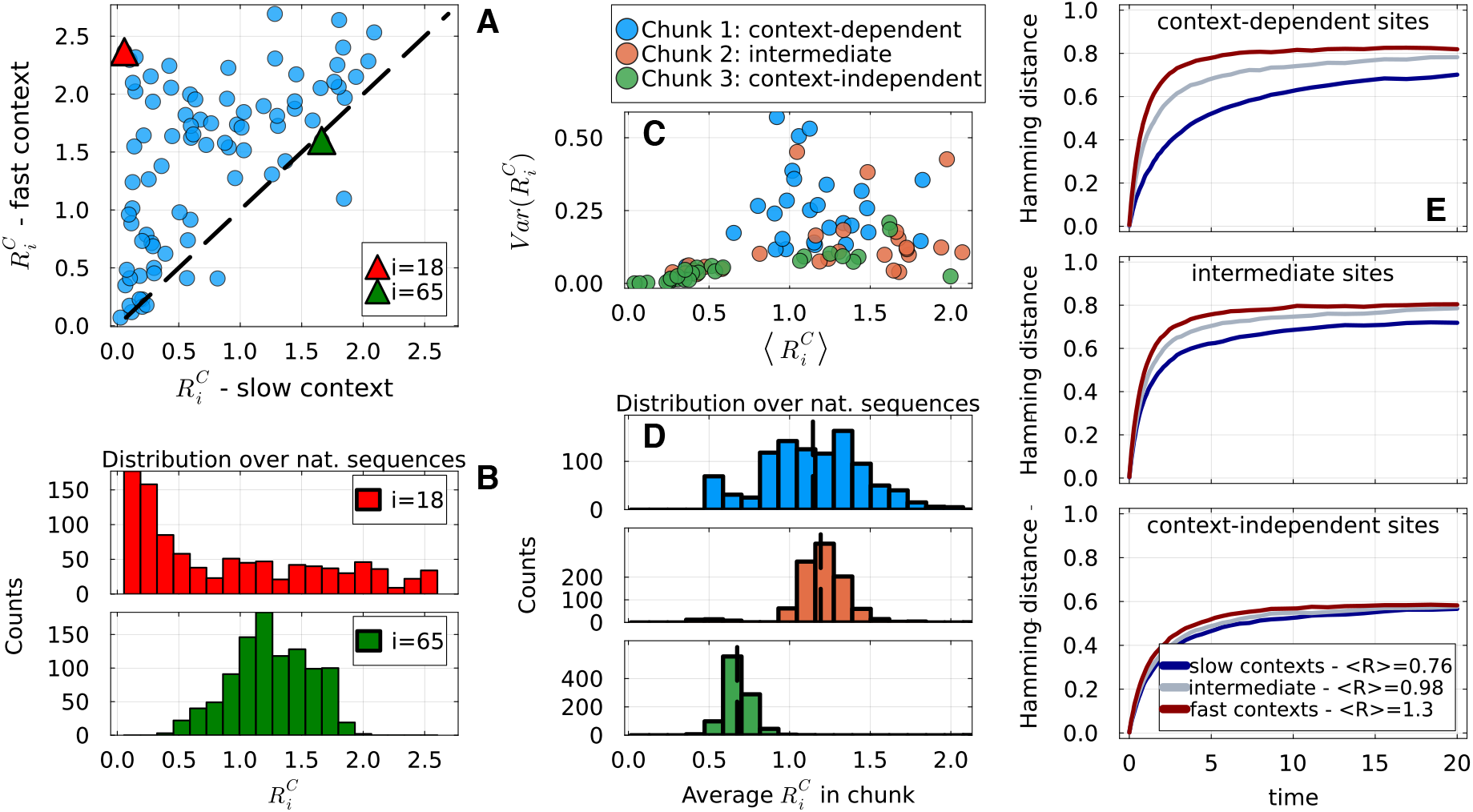
Equivalent to Figure 3 of the main text for the PF00595 family.

**Figure S14.**
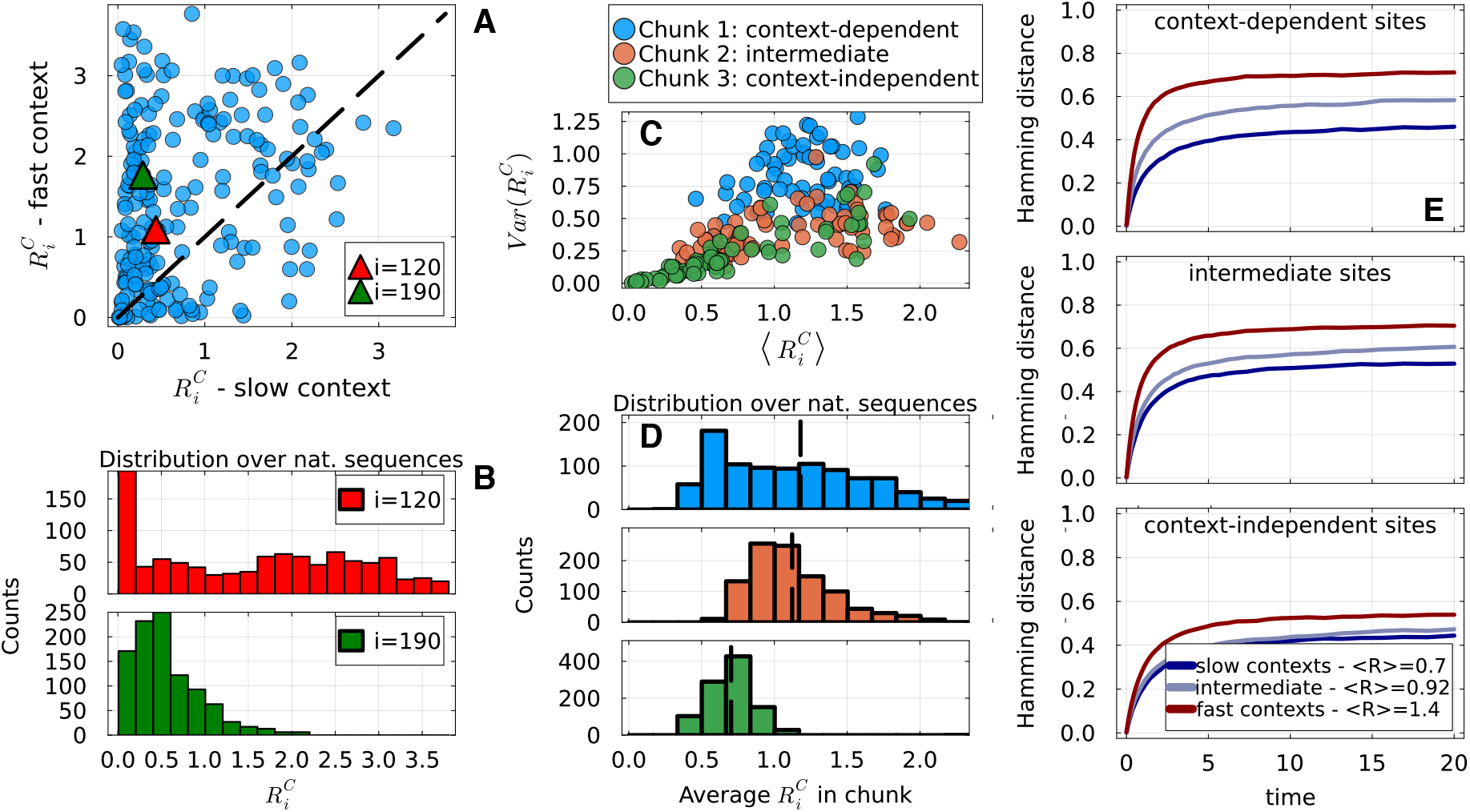
Equivalent to Figure 3 of the main text for the PF13354 family.

**Figure S15.**
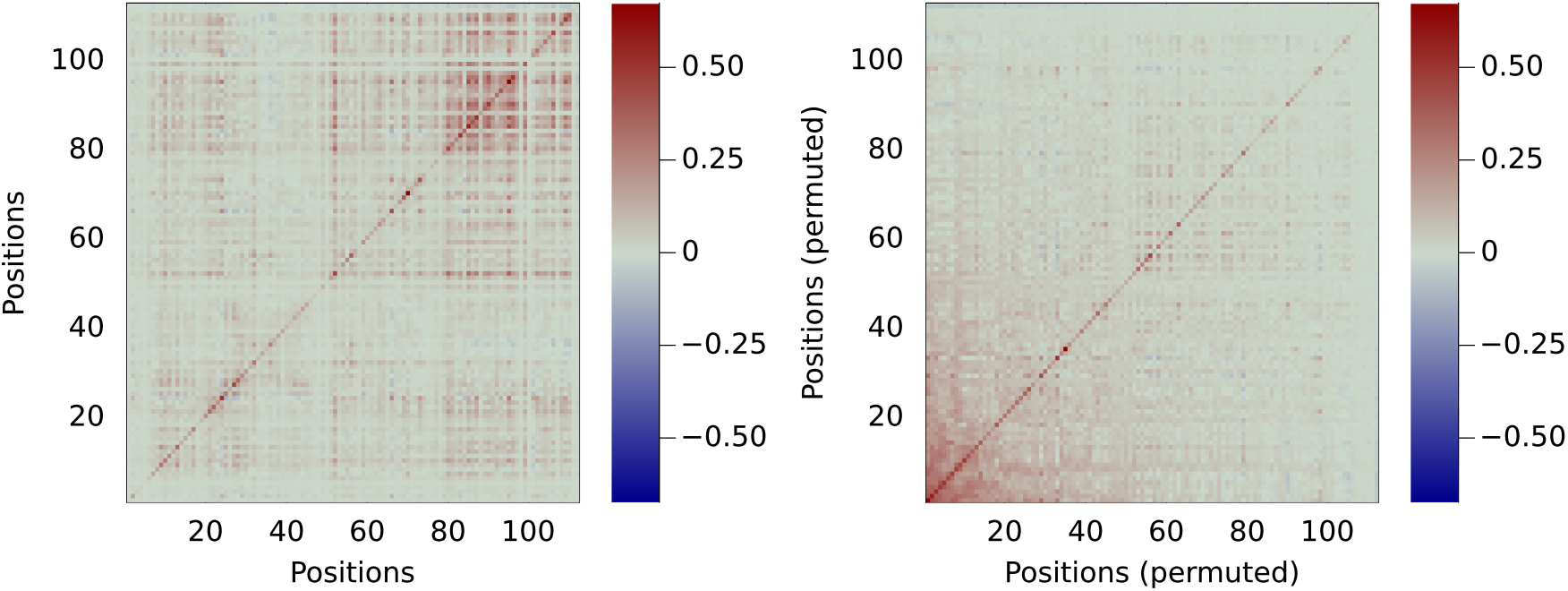
Heatmap of the covariance matrix of context-dependent rates: row *i* and column *j* corresponds to value 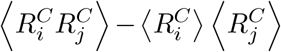, with the averages taken over sequences. **Left**: natural order of the positions. **Right**: Positions are permuted using the ranking procedure described in the main text. The ranking groups together positions that have a high or low context-dependent rate in the same contexts, *i*.*e*. high covariance.

**Figure S16.**
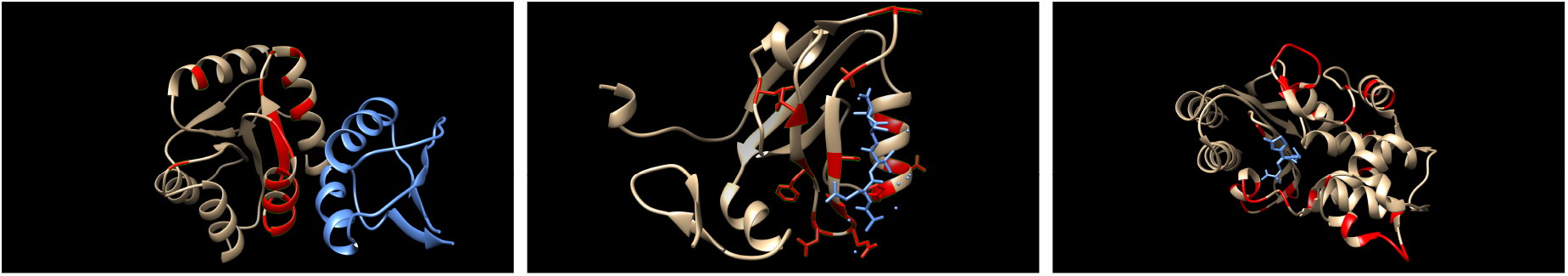
Structures of representents of the families used in the paper. We highlighted in red the *L/*5 sites with a strong context-dependence according to the procedure described around Figure 3, where *L* is the length of the sequence. **Left**: the CheY chemotaxis response-regulator domain (PF00072) of E. Coli (grey and red) in complex with its partner histidine kinease CheA (light blue) [57]. The top 22 context-dependent sites are structurally close and mostly located at the interface. PDB: 1EAY. **Center**: third PDZ domain of the PSD-95 protein (grey and red) in complex with its ligand CRIPT (light blue) [58]. The top 16 sites are structurally close and mostly located at the interface with the ligand. PDB: 1BE9. **Right**: TEM-1 beta-lactamase TEM-1 beta-lactamase (grey and red) with a covalent inhibitor bound in the active site (light blue). The top 39 context-dependent sites are highlighted in red. PDB: 1M40.

**Figure S17.**
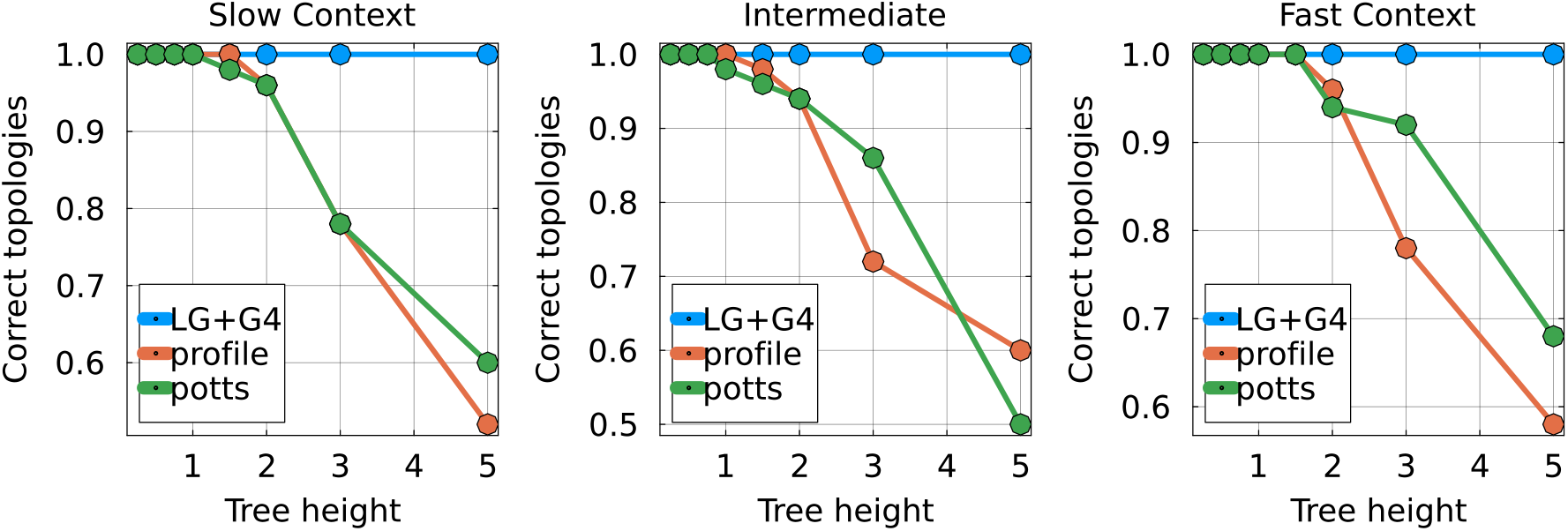
Fraction of correctly recovered topologies as a function of tree height in the four-taxon experiment, for different simulators. The correct tree has topology `((A,B),(C,D))` with all branches of equal length.

**Figure S18.**
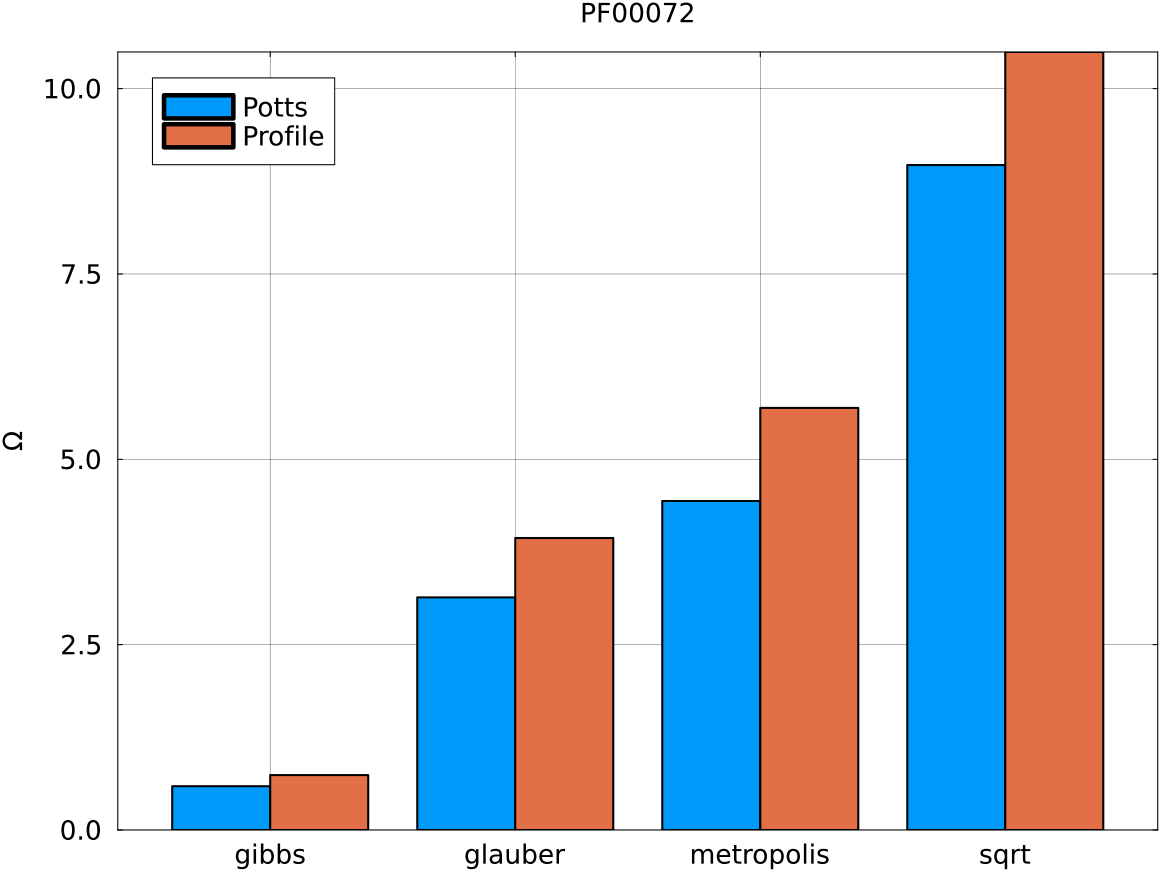
Value of the time normalization constant Ω for the PF00072 family and different evolutionary step types. Ω represents the amount of “physical” time that corresponds to one unit of evolutionary time.

**Figure S19.**
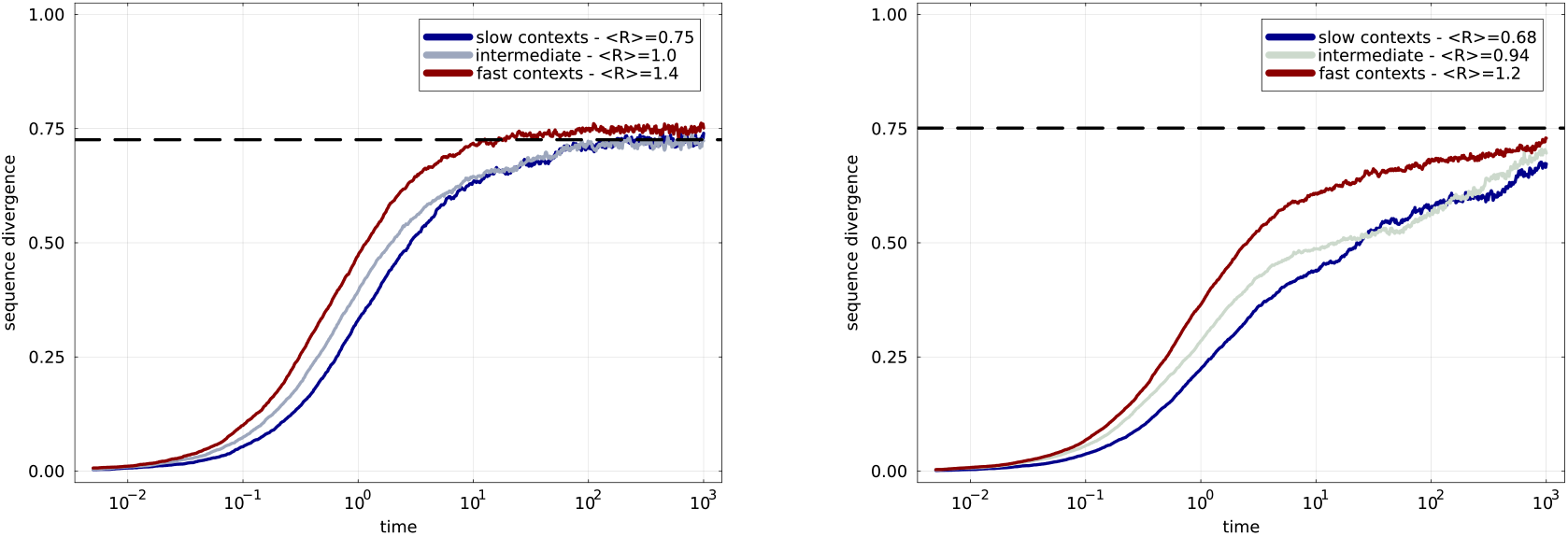
Similar to panel **E** of Figure 1 of the main text, but for longer evolutionary trajectories. Families are **left**: PF00072 and **right**: PF13354. Distances eventually saturate to a maximum even when starting from a slow context.

**Figure S20.**
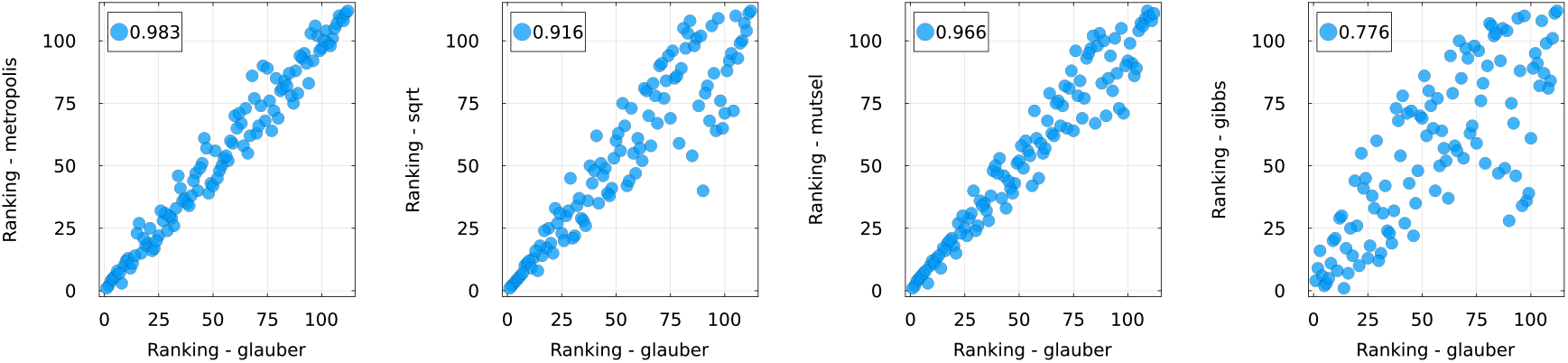
Ranking of positions based on their context-dependent rates for different dynamics. Each point represents a position, with its rank for the two dynamics displayed as *x* and *y* coordinates. The number indicates the Pearson correlation. Rankings are very similar for the Glauber, Metropolis, square-root and mutation-selection dynamics, especially for the top ranked sites. Gibbs dynamics are visibly different, although the ranking is still strongly correlated.

**Figure S21.**
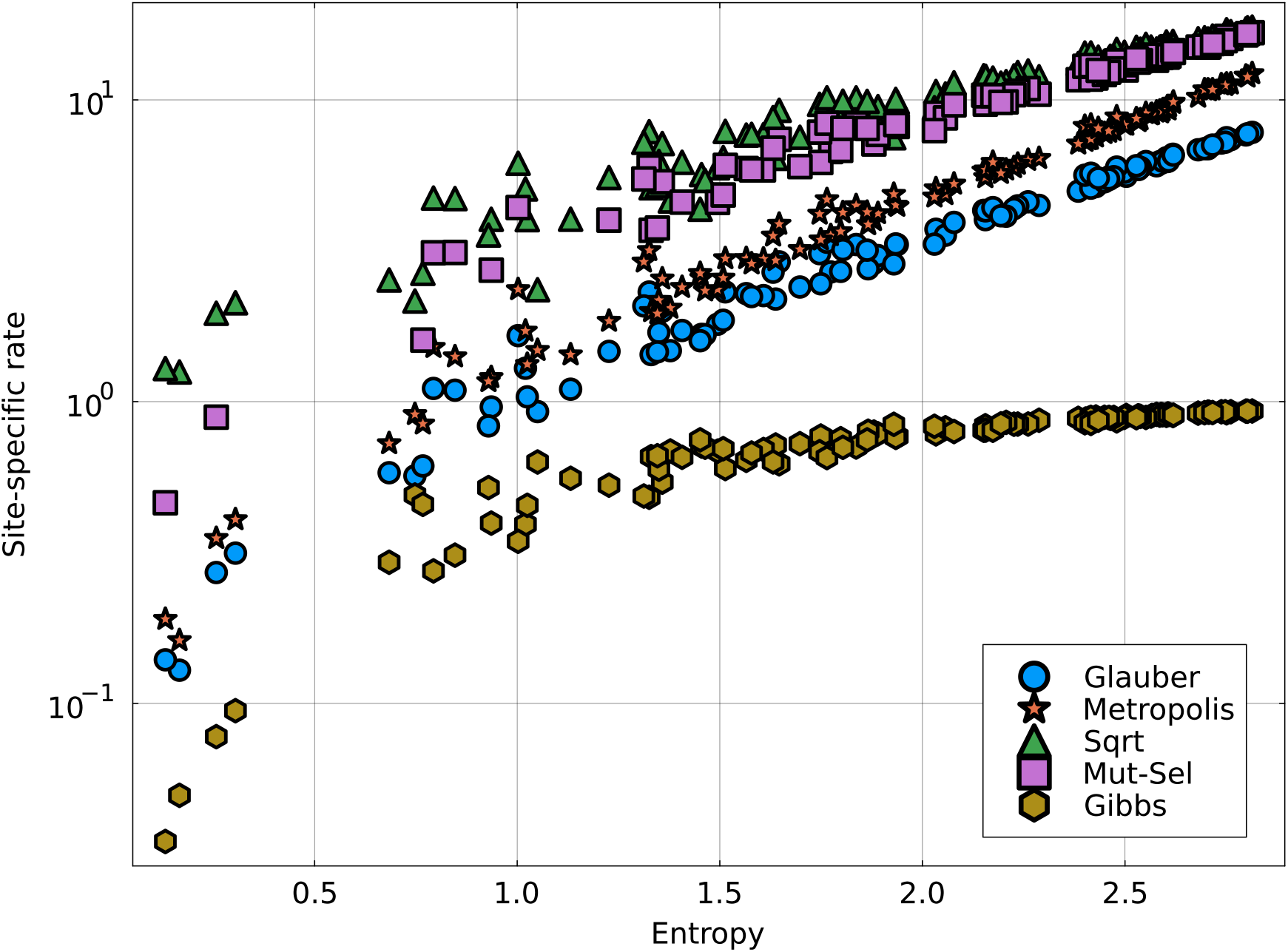
Analytical site-specific rates in the profile model (Eq. A10) against site entropy for all positions of the PF00072 alignment. The Gibbs rate is a probabilty and by construction saturates to one, which is not the case for the others.

## References

[1] Ziheng Yang. Maximum likelihood phylogenetic estimation from DNA sequences with variable rates over sites: Approximate methods. Journal of Molecular Evolution, 39(3):306–314, September 1994. ISSN 1432-1432. doi:10.1007/BF00160154.

[2] X Gu, Y X Fu, and W H Li. Maximum likelihood estimation of the heterogeneity of substitution rate among nucleotide sites. Molecular Biology and Evolution, 12(4):546–557, July 1995. ISSN 0737-4038. doi:10.1093/oxfordjournals.molbev.a040235.

[3] David T. Jones, William R. Taylor, and Janet M. Thornton. The rapid generation of mutation data matrices from protein sequences. Bioinformatics, 8(3):275–282, June 1992. ISSN 1367-4803. doi:10.1093/bioinformatics/8.3.275.

[4] Si Quang Le and Olivier Gascuel. An Improved General Amino Acid Replacement Matrix. Molecular Biology and Evolution, 25(7):1307–1320, July 2008. ISSN 0737-4038. doi: 10.1093/molbev/msn067.

[5] A L Halpern and W J Bruno. Evolutionary distances for protein-coding sequences: Modeling site-specific residue frequencies. Molecular Biology and Evolution, 15(7):910–917, July 1998. ISSN 0737-4038. doi:10.1093/oxfordjournals.molbev.a025995.

[6] Le Si Quang, Olivier Gascuel, and Nicolas Lartillot. Empirical profile mixture models for phylogenetic reconstruction. Bioinformatics, 24(20):2317–2323, October 2008. ISSN 1367-4803. doi:10.1093/bioinformatics/btn445.

[7] Nicolas Rodrigue, Hervé Philippe, and Nicolas Lartillot. Mutation-selection models of coding sequence evolution with site-heterogeneous amino acid fitness profiles. Proceedings of theNational Academy of Sciences, 107(10):4629–4634, March 2010. doi:10.1073/pnas.0910915107.

[8] Stephen M Crotty, Bui Quang Minh, Nigel G Bean, Barbara R Holland, Jonathan Tuke, Lars S Jermiin, and Arndt Von Haeseler. GHOST: Recovering Historical Signal from Heterotachously Evolved Sequence Alignments. Systematic Biology, 69(2):249–264, March 2020. ISSN 1063-5157.doi:10.1093/sysbio/syz051.

[9] Michael Socolich, Steve W. Lockless, William P. Russ, Heather Lee, Kevin H. Gardner, and Rama Ranganathan. Evolutionary information for specifying a protein fold. Nature, 437(7058): 512–518, September 2005. ISSN 1476-4687. doi:10.1038/nature03991.

[10] Tyler N. Starr and Joseph W. Thornton. Epistasis in protein evolution. Protein Science, 25 (7):1204–1218, 2016. ISSN 1469-896X. doi:10.1002/pro.2897.

[11] Johanna Trost, Julia Haag, Dimitri Höhler, Laurent Jacob, Alexandros Stamatakis, and Bastien Boussau. Simulations of Sequence Evolution: How (Un)realistic They Are and Why. Molecular Biology and Evolution, 41(1):msad277, January 2024. ISSN 1537-1719. doi:10.1093/molbev/msad277.

[12] William P. Russ, Matteo Figliuzzi, Christian Stocker, Pierre Barrat-Charlaix, Michael Socolich, Peter Kast, Donald Hilvert, Remi Monasson, Simona Cocco, Martin Weigt, and Rama Ranganathan. An evolution-based model for designing chorismate mutase enzymes. Science, 369(6502):440–445, July 2020. ISSN 0036-8075, 1095-9203. doi:10.1126/science.aba3304.

[13] Francisco McGee, Sandro Hauri, Quentin Novinger, Slobodan Vucetic, Ronald M. Levy,Vincenzo Carnevale, and Allan Haldane. The generative capacity of probabilistic protein sequence models. Nature Communications, 12(1):6302, November 2021. ISSN 2041-1723. doi:10.1038/s41467-021-26529-9.

[14] Jérôme Tubiana, Simona Cocco, and Rémi Monasson. Learning protein constitutive motifs from sequence data. eLife, 8:e39397, March 2019. ISSN 2050-084X. doi:10.7554/eLife.39397.

[15] Xinqiang Ding, Zhengting Zou, and Charles L. Brooks Iii. Deciphering protein evolution and fitness landscapes with latent space models. Nature Communications, 10(1):5644, December 2019. ISSN 2041-1723. doi:10.1038/s41467-019-13633-0.

[16] Roshan M. Rao, Jason Liu, Robert Verkuil, Joshua Meier, John Canny, Pieter Abbeel, Tom Sercu, and Alexander Rives. MSA Transformer. In Proceedings of the 38th International Conference on Machine Learning, pages 8844–8856. PMLR, July 2021.

[17] Alexander Rives, Joshua Meier, Tom Sercu, Siddharth Goyal, Zeming Lin, Jason Liu, Demi Guo, Myle Ott, C. Lawrence Zitnick, Jerry Ma, and Rob Fergus. Biological structure and function emerge from scaling unsupervised learning to 250 million protein sequences. Proceedings of the National Academy of Sciences, 118(15):e2016239118, April 2021. doi:10.1073/pnas.2016239118.

[18] Simona Cocco, Christoph Feinauer, Matteo Figliuzzi, Remi Monasson, and Martin Weigt. Inverse Statistical Physics of Protein Sequences: A Key Issues Review. Reports on Progress inPhysics, 81(3):032601, March 2018. ISSN 0034-4885, 1361-6633. doi:10.1088/1361-6633/aa9965.

[19] F. Morcos, A. Pagnani, B. Lunt, A. Bertolino, D. S. Marks, C. Sander, R. Zecchina, J. N. Onuchic, T. Hwa, and M. Weigt. Direct-coupling analysis of residue coevolution captures native contacts across many protein families. Proceedings of the National Academy of Sciences, 108 (49):E1293–E1301, December 2011. ISSN 0027-8424, 1091-6490. doi:10.1073/pnas.1111471108.

[20] Magnus Ekeberg, Cecilia Lövkvist, Yueheng Lan, Martin Weigt, and Erik Aurell. Improved contact prediction in proteins: Using pseudolikelihoods to infer Potts models. Physical Review E, 87(1), January 2013. ISSN 1539-3755, 1550-2376. doi:10.1103/PhysRevE.87.012707.

[21] Matteo Figliuzzi, Pierre Barrat-Charlaix, and Martin Weigt. How Pairwise Coevolutionary Models Capture the Collective Residue Variability in Proteins? Molecular Biology and Evolution, 35(4):1018–1027, April 2018. ISSN 0737-4038, 1537-1719. doi:10.1093/molbev/msy007.

[22] Linda Dib, Daniele Silvestro, and Nicolas Salamin. Evolutionary footprint of coevolving positions in genes. Bioinformatics, 30(9):1241–1249, May 2014. ISSN 1367-4803. doi: 10.1093/bioinformatics/btu012.

[23] Xavier Meyer, Linda Dib, Daniele Silvestro, and Nicolas Salamin. Simultaneous Bayesian inference of phylogeny and molecular coevolution. Proceedings of the National Academy of Sciences, 116(11):5027–5036, March 2019. doi:10.1073/pnas.1813836116.

[24] Andrew F Magee, Sarah K Hilton, and William S DeWitt. Robustness of Phylogenetic Inference to Model Misspecification Caused by Pairwise Epistasis. Molecular Biology and Evolution, 38 (10):4603–4615, October 2021. ISSN 1537-1719. doi:10.1093/molbev/msab163.

[25] Douglas M. Robinson, David T. Jones, Hirohisa Kishino, Nick Goldman, and Jeffrey L. Thorne. Protein Evolution with Dependence Among Codons Due to Tertiary Structure. Molecular Biology and Evolution, 20(10):1692–1704, October 2003. ISSN 0737-4038. doi: 10.1093/molbev/msg184.

[26] Nicolas Rodrigue HervéPhilippe, and Nicolas Lartillot. Assessing Site-Interdependent Phylogenetic Models of Sequence Evolution. Molecular Biology and Evolution, 23(9):1762–1775, September 2006. ISSN 0737-4038. doi:10.1093/molbev/msl041.

[27] Sanzo Miyazawa and Robert L. Jernigan. Estimation of effective interresidue contact energies from protein crystal structures: Quasi-chemical approximation. Macromolecules, 18(3):534–552, March 1985. ISSN 0024-9297. doi:10.1021/ma00145a039.

[28] Chris A. Nasrallah, David H. Mathews, and John P. Huelsenbeck. Quantifying the Impact of Dependent Evolution among Sites in Phylogenetic Inference. Systematic Biology, 60(1):60–73, January 2011. ISSN 1063-5157. doi:10.1093/sysbio/syq074.

[29] Jose Alberto de la Paz, Charisse M. Nartey, Monisha Yuvaraj, and Faruck Morcos. Epistatic contributions promote the unification of incompatible models of neutral molecular evolution. Proceedings of the National Academy of Sciences, page 201913071, March 2020. ISSN 0027-8424, 1091-6490. doi:10.1073/pnas.1913071117.

[30] David D. Pollock, Grant Thiltgen, and Richard A. Goldstein. Amino acid coevolution induces an evolutionary Stokes shift. Proceedings of the National Academy of Sciences, 109(21):E1352–E1359, May 2012. ISSN 0027-8424, 1091-6490. doi:10.1073/pnas.1120084109.

[31] Leonardo Di Bari, Matteo Bisardi, Sabrina Cotogno, Martin Weigt, and Francesco Zamponi. Emergent time scales of epistasis in protein evolution. Proceedings of the National Academy of Sciences, 121(40):e2406807121, October 2024. doi:10.1073/pnas.2406807121.

[32] Matteo Bisardi, Juan Rodriguez-Rivas, Francesco Zamponi, and Martin Weigt. Modeling Sequence-Space Exploration and Emergence of Epistatic Signals in Protein Evolution. Molecular Biology and Evolution, page msab321, November 2021. ISSN 1537-1719. doi:10.1093/molbev/msab321.

[33] Sophia Alvarez, Charisse M. Nartey, Nicholas Mercado, Jose Alberto de la Paz, Tea Huseinbegovic, and Faruck Morcos. In vivo functional phenotypes from a computational epistatic model of evolution. Proceedings of the National Academy of Sciences, 121(6):e2308895121, February 2024. doi:10.1073/pnas.2308895121.

[34] Indrani Choudhuri, Avik Biswas, Allan Haldane, and Ronald M. Levy. Contingency and Entrenchment of Drug-Resistance Mutations in HIV Viral Proteins. The Journal of Physical Chemistry B, 126(50):10622–10636, December 2022. ISSN 1520-6106. doi:10.1021/acs.jpcb.2c06123.

[35] Avik Biswas, Indrani Choudhuri, Eddy Arnold, Dmitry Lyumkis, Allan Haldane, and Ronald M. Levy. Kinetic coevolutionary models predict the temporal emergence of HIV-1 resistance mutations under drug selection pressure. Proceedings of the National Academy of Sciences, 121 (15):e2316662121, April 2024. doi:10.1073/pnas.2316662121.

[36] Daniel T. Gillespie. Exact stochastic simulation of coupled chemical reactions. The Journal of Physical Chemistry, 81(25):2340–2361, December 1977. ISSN 0022-3654. doi: 10.1021/j100540a008.

[37] Jaina Mistry, Sara Chuguransky, Lowri Williams, Matloob Qureshi, Gustavo A Salazar, Erik L L Sonnhammer, Silvio C E Tosatto, Lisanna Paladin, Shriya Raj, Lorna J Richardson, Robert DFinn, and Alex Bateman. Pfam: The protein families database in 2021. Nucleic Acids Research,page gkaa913, October 2020. ISSN 0305-1048, 1362-4962. doi:10.1093/nar/gkaa913.

[38] Anna Paola Muntoni, Andrea Pagnani, Martin Weigt, and Francesco Zamponi. adabmDCA: Adaptive Boltzmann machine learning for biological sequences. BMC Bioinformatics, 22(1):528, October 2021. ISSN 1471-2105. doi:10.1186/s12859-021-04441-9.

[39] Nhan Ly-Trong, Suha Naser-Khdour, Robert Lanfear, and Bui Quang Minh. AliSim: A Fastand Versatile Phylogenetic Sequence Simulator for the Genomic Era. Molecular Biology and Evolution, 39(5):msac092, May 2022. ISSN 1537-1719. doi:10.1093/molbev/msac092.

[40] P. Lopez, D. Casane, and H. Philippe. Heterotachy, an Important Process of Protein Evolution. Molecular Biology and Evolution, 19(1):1–7, January 2002. ISSN 0737-4038. doi: 10.1093/oxfordjournals.molbev.a003973.

[41] Julien Soubrier, Mike Steel, Michael S.Y. Lee, Clio Der Sarkissian, Stéphane Guindon, Si-mon Y.W. Ho, and Alan Cooper. The Influence of Rate Heterogeneity among Sites on the Time Dependence of Molecular Rates. Molecular Biology and Evolution, 29(11):3345–3358, November 2012. ISSN 0737-4038. doi:10.1093/molbev/mss140.

[42] Bui Quang Minh, Heiko A Schmidt, Olga Chernomor, Dominik Schrempf, Michael D Woodhams, Arndt von Haeseler, and Robert Lanfear. IQ-TREE 2: New Models and Efficient Methods for Phylogenetic Inference in the Genomic Era. Molecular Biology and Evolution, 37(5):1530–1534, May 2020. ISSN 0737-4038. doi:10.1093/molbev/msaa015.

[43] Subha Kalyaanamoorthy, Bui Quang Minh, Thomas K. F. Wong, Arndt von Haeseler, and Lars S. Jermiin. ModelFinder: Fast model selection for accurate phylogenetic estimates. Nature Methods, 14(6):587–589, June 2017. ISSN 1548-7105. doi:10.1038/nmeth.4285.

[44] Luca Nesterenko, Luc Blassel, Philippe Veber, Bastien Boussau, and Laurent Jacob. Phyloformer: Fast, Accurate, and Versatile Phylogenetic Reconstruction with Deep Neural Net-works. Molecular Biology and Evolution, 42(4):msaf051, April 2025. ISSN 1537-1719. doi:10.1093/molbev/msaf051.

[45] Anne-Florence Bitbol, Robert S. Dwyer, Lucy J. Colwell, and Ned S. Wingreen. Inferring interaction partners from protein sequences. Proceedings of the National Academy of Sciences of the United States of America, 113(43):12180–12185, October 2016. ISSN 0027-8424. doi: 10.1073/pnas.1606762113.

[46] Matteo Figliuzzi, Hervé Jacquier, Alexander Schug, Oliver Tenaillon, and Martin Weigt. Coevolutionary Landscape Inference and the Context-Dependence of Mutations in Beta-Lactamase TEM-1. Molecular Biology and Evolution, 33(1):268–280, January 2016. ISSN 0737-4038, 1537-1719. doi:10.1093/molbev/msv211.

[47] Simona Cocco, Lorenzo Posani, and Rémi Monasson. Functional effects of mutations in proteins can be predicted and interpreted by guided selection of sequence covariation information.Proceedings of the National Academy of Sciences, 121(26):e2312335121, June 2024. doi:10.1073/pnas.2312335121.

[48] Guy Sella and Aaron E. Hirsh. The application of statistical physics to evolutionary biology. Proceedings of the National Academy of Sciences, 102(27):9541–9546, July 2005. doi: 10.1073/pnas.0501865102.

[49] Christoph Feinauer, Marcin J. Skwark, Andrea Pagnani, and Erik Aurell. Improving Contact Prediction along Three Dimensions. PLOS Computational Biology, 10(10):e1003847, October 2014. ISSN 1553-7358. doi:10.1371/journal.pcbi.1003847.

[50] Matteo De Leonardis, Andrea Pagnani, and Pierre Barrat-Charlaix. Reconstruction of Ancestral Protein Sequences Using Autoregressive Generative Models. Molecular Biology and Evolution, 42(4):msaf070, April 2025. ISSN 1537-1719. doi:10.1093/molbev/msaf070.

[51] Jeanne Trinquier, Guido Uguzzoni, Andrea Pagnani, Francesco Zamponi, and Martin Weigt.Efficient generative modeling of protein sequences using simple autoregressive models. Nature Communications, 12(1):5800, October 2021. ISSN 2041-1723. doi:10.1038/s41467-021-25756-4.

[52] R. D. Finn, J. Clements, and S. R. Eddy. HMMER web server: Interactive sequence similarity searching. Nucleic Acids Research, 39(suppl):W29–W37, July 2011. ISSN 0305-1048, 1362-4962. doi:10.1093/nar/gkr367.

[53] The UniProt Consortium. UniProt: The universal protein knowledgebase in 2021. Nucleic Acids Research, 49(D1):D480–D489, January 2021. ISSN 0305-1048. doi:10.1093/nar/gkaa1100.

[54] Ciyou Zhu, Richard H. Byrd, Peihuang Lu, and Jorge Nocedal. Algorithm 778: L-BFGS-B: Fortran subroutines for large-scale bound-constrained optimization. ACM Trans. Math. Softw., 23(4):550–560, December 1997. ISSN 0098-3500. doi:10.1145/279232.279236.

[55] Jarrett Revels, Miles Lubin, and Theodore Papamarkou. Forward-Mode Automatic Differentiation in Julia, July 2016.

[56] Motoo Kimura. Diffusion models in population genetics. Journal of Applied Probability, 1(2):177–232, December 1964. ISSN 0021-9002, 1475-6072. doi:10.2307/3211856.

[57] Megan M. McEvoy, Andrew C. Hausrath, Gannon B. Randolph, S. James Remington, andFrederick W. Dahlquist. Two binding modes reveal flexibility in kinase/response regulator interactions in the bacterial chemotaxis pathway. Proceedings of the National Academy of Sciences, 95(13):7333–7338, June 1998. doi:10.1073/pnas.95.13.7333.

[58] Declan A. Doyle, Alice Lee, John Lewis, Eunjoon Kim, Morgan Sheng, and Roderick MacKinnon.Crystal Structures of a Complexed and Peptide-Free Membrane Protein–Binding Domain: Molecular Basis of Peptide Recognition by PDZ. Cell, 85(7):1067–1076, June 1996. ISSN 0092-8674, 1097-4172. doi:10.1016/S0092-8674(00)81307-0.

